# TLR3 Deficiency Leads to Altered Immune Responses to *Chlamydia trachomatis* Infection in Human Oviduct Epithelial Cells

**DOI:** 10.1101/680793

**Authors:** Jerry Z. Xu, Ramesh Kumar, Haoli Gong, Luyao Liu, Nicole Ramos-Solis, Yujing Li, Wilbert A. Derbigny

**Author notes:** These authors contributed equally to this work (Jerry Z. Xu, Ramesh Kumar).

## Abstract

Reproductive tract pathology caused by *Chlamydia trachomatis* infection is an important global cause of human infertility. To better understand the mechanisms associated with *Chlamydia*-induced genital tract pathogenesis in humans, we used CRISPR genome editing to disrupt TLR3 function in the human oviduct epithelial (hOE) cell-line OE-E6/E7, in order to investigate the possible role(s) of TLR3 signaling in the immune response to *Chlamydia*. Disruption of TLR3 function in these cells significantly diminished the *Chlamydia*-induced synthesis of several inflammation biomarkers including IFN-β, IL-6, IL-6Ra, sIL-6Rβ (gp130), IL-8, IL-20, IL-26, IL-34, sTNF-R1, TNFSF13B, MMP-1, MMP-2, and MMP-3. In contrast, the *Chlamydia*-induced synthesis of CCL-5, IL-29 (IFNλ1) and IL-28A (IFNλ2) were significantly *increased* in the TLR3-deficient hOE cells when compared to their wild-type counterparts. Our results propose a role for TLR3 signaling in limiting the genital tract fibrosis, scarring, and chronic inflammation often associated with human chlamydial disease. Interestingly, we saw that *Chlamydia* infection induced the production of biomarkers associated with persistence, tumor metastasis, and autoimmunity such as soluble CD163 (sCD163), chitinase-3-like protein 1, osteopontin, and pentraxin-3 in the hOE cells; however, their expression levels were significantly dysregulated in the TLR3-deficient hOE cells. Finally, we demonstrate using the hOE cells that TLR3 deficiency resulted in an increased amount of chlamydial LPS within the *Chlamydia* inclusion, which is suggestive that TLR3 deficiency leads to enhanced chlamydial replication and possibly increased genital tract pathogenesis during human infection.

**Abbreviations:** hOE, human OE-E6/E7 cells; TLR3 KO, TLR3 knockout cell line; poly (I:C), Polyinosinic–polycytidylic acid sodium salt.

## Introduction

The bacterial pathogen *Chlamydia trachomatis* has caused 1,526,658 infections in the United States in 2015 (an increase of 6% since 2014), and it is the most commonly reported bacterial sexually transmitted disease (STD) in the United States (1). Genital tract *C. trachomatis* infections are easily cured with antibiotics if properly diagnosed at early stages of infection. However, because 75-90% of women infected with *Chlamydia* are asymptomatic for clinical disease, opportunities for therapeutic interventions are usually missed. The asymptomatic nature of the clinical symptoms is the major contributing factor to the continuing spread of the disease to uninfected partners, and the more severe pathogenesis and sequelae that often lead to infertility in women. Further contributing to the growing rates of infectivity amongst the previously uninfected populations are the statistics showing that up to 90% of men infected with *Chlamydia* exhibit no symptoms (2, 3) and that an effective vaccine remains elusive (4).

*Chlamydia* infections are also leading causes of pelvic inflammatory disease (5), tubal occlusion (6), and ectopic pregnancy (7, 8) in women. Interactions between the host immunity and *Chlamydia* infection are thought to be largely responsible for the pathology associated with human chlamydial disease, though the precise pathogenic mechanisms remain unclear (9, 10). As an obligate intracellular pathogen, *Chlamydia* species are known to interact with host-cell pattern recognition receptors (PRRs), including a variety of intracellular cytosolic receptors and Toll-like receptors (TLRs), to trigger the innate-immune inflammatory response (11–18). Stimulation of genital tract epithelial cell TLRs (and other PRRs) by chlamydial pathogen-associated molecular patterns (PAMPs) trigger cytokine responses that are critical to the establishment of innate and adaptive immunity. These *Chlamydia*-induced cytokine responses also include the syntheses of inflammatory mediators that have been implicated as the major culprit in the pathology associated with *Chlamydia* disease (12, 14, 19–24). The overall goal of these investigations into the interactions between host-cell PRRs and *Chlamydia* infection is to identify the PRRs that trigger specific inflammatory mediators that cause scarring and fibrosis, and then define therapeutic measures to prevent that process.

Human genital tract epithelial cells express most of the known TLRs; however, the TLRs are known to vary in their expression levels within the female reproductive tract (depending upon the concentration of specific sex hormones) and their tissue distribution (25). The human Fallopian-derived epithelial cell line OE-E6/E7 (26) was shown to express functional proteins for TLRs 1 through 6, of which TLR2 was shown to have a role in the innate-immune response to *C. trachomatis* infection (27, 28). TLR2 has also been shown to have a role in the immune responses to *C. muridarum* infection in mice, and that it had a significant role in the *Chlamydia*-induced genital tract pathology observed in the infected animals (12, 29–31). In our early investigations into the role of TLR3 in the immune response to genital tract *Chlamydia* infection, we showed that *C. muridarum* infected murine oviduct epithelial (OE) cells secrete IFN-β in a mostly TLR3 dependent manner, and that mouse OE cells deficient in TLR3 show dramatic reductions in the synthesis of other inflammatory immune mediators in addition to IFN-β (13, 15). Our most recent report shows that TLR3-deficient mice have increased chlamydial loads, aberrant genital tract secretion levels of several different inflammatory mediators, an altered CD4^+^ T-cell recruitment, and more severe oviduct and uterine horn pathology when compared to wild-type controls (32). Our data propose a protective role for TLR3 signaling in the immune response to *Chlamydia* infection in mice. However, the role of TLR3 in the immune response to *Chlamydia* infection in human oviduct tissue has not yet been investigated and remains unclear. In this study, we used the immortalized human oviduct epithelial cell line OE-E6/E7 to assess the role of TLR3 in the immune responses to *Chlamydia trachomatis* L2 infection.

## MATERIALS AND METHODS

### Cell culture

Human OE-E6/E7 (hOE) cells and Hela cells were maintained in high glucose Dulbecco’s modified Eagle’s medium (Life Technologies, Inc.) supplemented with 10% bovine calf serum (Hyclone) in a 37 °C, 5% CO_2_ incubator. The hOE cells were originally derived from Fallopian tube tissue and were immortalized by expressing the HPV 16 E6/E7 open reading frame in a retroviral expression system (26).

### Reagents

The following TLR agonists were purchased from InVivoGen (San Diego, CA): 1) peptidoglycan from *E. coli* serotype 0111:B4 (125 EU/mg); 2) ultrapure flagellin purified from isolated from *Bacillus subtilis* (>95% purity); and 3) ODN 2216, a synthetic oligonucleotide (ODN) that preferentially binds human TLR9 and ODN2243, an ODN 2216 control without CpGs. Wildtype (WT) and TLR3-deficient hOE cells were grown to confluence in 24-well before being treated with the appropriate TLR ligand at the concentrations specified in the text. Supernatants were harvested at the 24h post-treatment time point and analyzed for cytokine content using ELISA for IL-6 (R&D Systems; Minneapolis, MN) according to the manufacturer’s protocol

### Generating theTLR3-deficient human epithelial cell lines

The human OE-E6/E7 cells and Hela cells were grown to 60-70% confluence before being transduced with either the human TLR3 CRISPR knockout Lentivirus or the CRISPR-Lenti non-targeting control transduction particles and 4µg/µL Polybrene. The Lenti-CRISPR Transduction particles (pLV-U6g-EPCG-TLR3), All-in-one ready-to-use Cas9 and guide RNA (gRNA), and CRISPR negative controls were purchased from Sigma (Sigma-Aldrich; St Louis, MO). The CRISPR system consisted of 3 gRNA sequences (CCACCTGAAGTTGACTCAGGTA, CCAACTTCACAAGGTATAGCCA, and CAGGGTGTTTTCACGCAATTGG), which targeted the human TLR3 gene (NM_003265) at exon 2, exon 2, and exon 4 respectively. The CRISPR Universal negative control targets no known human, mouse, or rat gene. The transduced human OE-E6/E7 cells and Hela cells were subjected to 5µg/mL puromycin selection, and the puromycin-resistant cells were sorted by using BD FACSAria cell sorter. The brightest GFP positive cells were collected and cultured for further single-cell cloning by glass cloning cylinders. Individual cell colonies were isolated and expanded. Higher GFP expressing cell clones were further selected by using BD Accuri C6 flow cytometer. Confirmation of TLR3 gene deletion, protein expression, and receptor function was assessed using PCR, western blot, and ELISA, respectively.

### Genomic DNA purification and PCR-amplification

Genomic DNA (gDNA) from hOE-TLRKO was purified by using the PureLink Genomic DNA Mini Kit from Invitrogen (Invitrogen Catlog# K1820-01). The gene-specific primer pairs (forward 5’-ACA AGG AAT ATA CCA ATG CAT TTG-3’and reverse 5’-GAT ATT TAG ATA GTA AGT CTA AGG-3’) were designed approximate 300 bps Up-and Down streams of the gRNA (CCACCTGAAGTTGACTCAGGTA) sequence spanning exon-2 of TLR3 gene. The purified g-DNA as a template and Platinum Super-Fi PCR master mix (Invitrogen Catlog# 12358-010) were used to amplify the PCR product (599bps). The cycling conditions were as: initial denaturation at 95°C for 2 min followed by 32 cycles, denaturation at 95°C for 45 s, annealing at 50.5°C for 45 s, elongation at 72°C for 45 s and final elongation at 72°C for 2 min. The amplified PCR products were gel purified using the QIAquick gel extraction kit (Qiagen; Germantown, MD). The PCR product from hOE-WT cells was used as control sample throughout this experiment. A portion of the amplified PCR products was sent for sequencing at the IUSM Bioinformatics Core Sequencing Facility, while the rest of the PCR products were used in the cloning experiment.

### Cloning and Plasmid Purification

An additional adenine-overhang was added to the portion of the PCR product to be cloned by using hi-fidelity Taq-polymerase and dATP at 70°C for 20min. The reaction mixtures were then ligated into pGEM-T Easy vector (Promega Catlog#A1360) overnight at 4°C and then transformed into TOP10 competent *E. coli*. The transformed *E. coli* were plated on LB-agar plates containing ampicillin (60ug/ml) supplemented with IPTG. The blue and white colonies were screened and inoculated into 5ml LB medium containing Amp (final concentration 60ug/ml). The putative positive cultures were used for plasmid purification by using QiaSpin Miniprep Kit (Catlog# 27106), and inserts were confirmed by PCR before plasmid sequencing.

### Chlamydia trachomatis preparation

The *C. trachomatis* L2-434/Bu (L2) and *C. trachomatis* UW-3CX serovar D strains were generously provided by Dr. David E. Nelson. The *C. trachomatis*-serovar D and L2 mother pools used in these experiments were subsequently grown and titrated as described in (33), whereby antibody specific for chlamydial LPS was used to identify chlamydial inclusions in the infected Hela cells. Alexa Fluor 488 and Alexa Fluor 594 anti-mouse IgG antibodies used in these experiments were purchased from Invitrogen (Invitrogen/Life Technologies; Carlsbad, CA), and immuno-staining results for titration of infectious chlamydial elementary bodies (EBs) were scanned and recorded by EVOS FL auto cell imaging system (Thermo-Fisher, Pittsburgh, PA). The corresponding isotype controls were used as negative controls.

### hOE and Hela cell Infections

OE-E6/E7 and Hela cells were plated in either 12 or 6-well tissue culture plates and grown to 80-90% confluence. For all experiments (unless stated otherwise), the cells were infected with 10 inclusion-forming-units (IFU) of *C. trachomatis*/ cell in cell culture medium. Immediately after adding *Chlamydia*, the tissue culture plates were gently agitated, centrifuged at 1200 rpm (200 × g) in a table-top centrifuge for 1 hour, and then incubated at 37°C in a 5% CO_2_ humidified incubator without subsequent change of medium until the time of cell harvest. Mock-infected control cells received an equivalent volume of epithelial cell culture medium but lacked any viable *C. trachomatis*.

### Protein extraction and evaluating protein expression levels

Epithelial cell lysates were prepared via incubation of PBS washed cell monolayers in RIPA buffer (EMD Millipore; Burlington, Massachusetts) with the addition of 1mM PMSF and 1x Protease Inhibitor Cocktails (Sigma). The total protein concentration of each sample was determined by using the Pierce BCA protein assay (Thermo Scientific). Analyses of protein expression were performed by using the WES™ simple western system (ProteinSimple; San Jose, California). Endogenous TLR3 protein expression in the wild type (WT) and CRISPR-modified epithelial cells was detected using a 1:50 dilution of TLR3 monoclonal antibody (Abcam; Cambridge, MA). Relative TLR3 protein expression levels between WT and CRISPR modified cells were obtained by measuring TLR3’s expression against the intracellular β-Actin control protein bound to an anti-β-Actin monoclonal antibody (1:300; Sigma). Protein detection in WES was accomplished according to the manufacturer’s protocol using streptavidin-HRP based methodology (ProteinSimple). The positive control for testing antibody specificity for TLR3 expression included loading (in separate reactions) either 50ng or 100ng of HEK293 cell lysate overexpressing human TLR3 (Novus).

### RNA purification and real-time quantitative PCR (RT-qPCR)

Total cell RNA was isolated from mock and *C. trachomatis*-infected WT and TLR3-deficient epithelial cells using the RNeasy kit plus (Qiagen, Valencia, CA). The DNA-free RNA was quantified using the NanoDrop spectrophotometer (Thermo Scientific), and cDNA was obtained with the Applied Biosystem’s high-capacity cDNA reverse transcription kit (Thermo Fisher). Target mRNA was quantified using Applied Biosystem’s TaqMan gene-expression master kit in reactions containing either human TLR3 primers, human IFN-β primers, and/or β-actin control primers. Quantitative measurements were performed via the ABI7500 real-time PCR detection system (Thermo Fisher).

### Standard and Multiplex ELISA analyses

The human IFN-β ELISA Kit (cat. #41410-1) and IL-6 ELISA kit (cat. #D6050) purchased from R&D systems were used to measure the *Chlamydia*-induced IFN-β and IL-6 respectively, using the manufacturer’s protocol. The standard ELISA kits were used to measure single cytokines secreted into the supernatants of hOE-WT, hOE-N-Ctrl, and hOE-TLRKO-cells that were either mock treated, infected with *C. trachomatis*, or treated with various TLR agonist.

To measure the chlamydial induced syntheses of several immune factors simultaneously, WT and TLR3 deficient hOE cells were either mock treated or infected at a MOI of 10 IFU/ cell with either serovar D or the L2 strain of *C. trachomatis*. For *C. trachomatis*-L2 infections, the supernatants were harvested at 0 (mock), 6, 12, 18, 24, and 30hrs post infection. The supernatants from the L2 infections were subjected to multiplex ELISA analyses in triplicate setting using the Bio-Plex Pro human inflammation panel-1, 37-Plex #171AL001M (Bio-Rad; Hercules, California) according to the manufacturer’s instructions. For infections with *C. trachomatis*-serovar D, the supernatants were harvested at 0 (mock), 6, 12, 24, 36, 48, 60, and 72hrs post infection. The supernatants from the serovar D infections were subjected to multiplex ELISA analyses in triplicate setting using a custom designed 27-plex human magnetic Luminex assay #LXSAHM (R&D Systems) according to the manufacturer’s instructions. Analyses of the data were performed in concert with the Indiana University Multiplex Analysis Core located in the Melvin and Bren Simon Cancer Center.

### RNA-interference

The transfections of the TLR3-specific siRNA (cat. #AM16708; Ambion/ Thermo-Fisher) and the scrambled control RNA (Silencer™ negative control; Thermo-Fisher) were done using Lipofectamine RNAiMAX (Thermo-Fisher) as described in (34). Briefly, 75-80% confluent hOE-WT cells were transfected with 2.5ug of each siRNA for 24 hours at 37°C with 5% CO2. After the 24h period, cell supernatants were replaced with fresh media prior to being infected with 10 IFU/cell *C. trachomatis*-L2. The level of IFN-β expression was determined at the specified time post infection by ELISA as described above.

### Flow cytometric analysis and multi-spectral imaging flow cytometry

WT and TLR3 deficient epithelial cells were either mock treated or infected at a MOI of 10 IFU/ cell with *Chlamydia trachomatis* L2 for 24 hours. At the 24hr time point, monolayers were washed once with PBS/2mM EDTA before being gently removed from the plate with trypsin-versene, washed 3 times in ice-cold PBS/EDTA, and cell pellets resuspended in 4% Formalin for 30 min at room temperature. The cells were washed 3 times with PBS/ EDTA and were permeabilized in 0.3% Triton X-100/ PBS/EDTA for 5 min at room temperature. The cells were blocked in blocking buffer (5% FBS/0.1%BSA/0.1%Triton X-100/PBS/EDTA) at room temperature for 60min. The cells were stained with anti-chlamydia LPS antibody (from Dr. David E. Nelson) in blocking buffer (1:5) for 60 min at room temperature. The cells were washed 3 times with PBS/EDTA. The cells were further stained with secondary antibody APC anti-mouse IgG(H+L) (1:1000, ThermoFisher) in blocking buffer for 30min before being washed 3 times with PBS/EDTA. Finally, the cells were suspended in 0.6ml 2% FBS/PBS/EDTA for flow cytometry analysis. Cellular responses to *Chlamydia* infection were analyzed via BD LSRFortessa cell analyzer (Becton Dickinson; Franklin Lakes, NJ) or by Multi-spectral Imaging Flow Cytometer Amnis Image StreamX MKII (EMD Millipore). Data were interpreted by using FlowJo v10 (FlowJo, LLC) and IDEAS (EMD Millipore) software.

### Immunofluorescent Staining

WT and TLR3 deficient hOE cells were seeded in 96-well <u>µ-plate (Ibidi; Fitchburg, WI)</u> and allowed to grow to 90% confluence before being either mock treated or infected at a MOI of 5 IFU/ cell with *C. trachomatis*-serovar D. At 36 hours post infection, the infected cells were then fixed with 200 µl of methanol and incubate at room temperature for 10 minutes. The fixed cells were stained with a 1:100 dilution of anti-chlamydial LPS antibody (EVI-HI; provided to us by Dr. David E. Nelson) and incubated for 1 hour at room temperature. The stained cells were washed 3 times with PBS. Detection was accomplished with a secondary stain of Alexa Fluor 488 and Alexa Fluor 594 anti-mouse IgG antibodies (Invitrogen). Nuclei were counterstained with 4,6-diamidino-2-phenylindole (DAPI; Life Technologies) according to manufacturer’s protocol and imaged at 60X under oil immersion using a Nikon Eclipse Ti microscope.

### Statistical Analysis

Numerical data are presented as means ± SEM. All experiments were repeated at least three times, and statistical significance was determined using either 2-way *ANOVA* analyses in GraphPad Prism or Student’s two-tailed *t*-test. Statistically significant differences are shown by asterisks (*) and with the minimum criteria being p <0.05.

## RESULTS

### IFN-β is induced in human OE-E6/7 cells in response to *Chlamydia* infection

IFN-β is known to be expressed during activation of the TLR3 signaling pathway during certain viral infections, and by stimulation via the synthetic double-stranded RNA analog poly (I:C) (35, 36). To confirm the presence of TLR3 and ascertain its function in the human OE-E6/E7 cells (hOE), the hOE cells were incubated in cell culture media supplemented with increasing concentrations of poly (I:C) for 24hrs. Figure 1A shows that the relative IFN-β mRNA expression was increased at the concentrations of 25, 50, and 100 µg/mL when compared to untreated controls. These results confirm that the TLR3 is functional in the hOE cells by demonstrating a dose-dependent increase in IFN-β gene expression in response to poly (I:C) stimulation. To ascertain the impact of *Chlamydia* infection on IFN-β synthesis in the hOE cells, we next infected hOE cells with *Chlamydia trachomatis* L2 at a MOI of 10 IFU/ cell for 24hrs, and measured the mRNA expression levels of both IFN-β and TLR3. As shown in Figures 1B and 1C, mRNA expression levels of IFN-β and TLR3 were increased during *Chlamydia* infection. These data are suggestive that *Chlamydia* infection of the hOE cells induces IFN-β synthesis and upregulates TLR3 gene expression in human oviduct tissue in a similar manner to what we have observed in the murine oviduct epithelial cells (14, 24). These findings provide the impetus for us to extrapolate that the IFN-β induced during *Chlamydia* infection in hOE cells will also occur via TLR3-dependent mechanisms similar to what we have reported in the murine OE cells (13).

**FIGURE 1.**
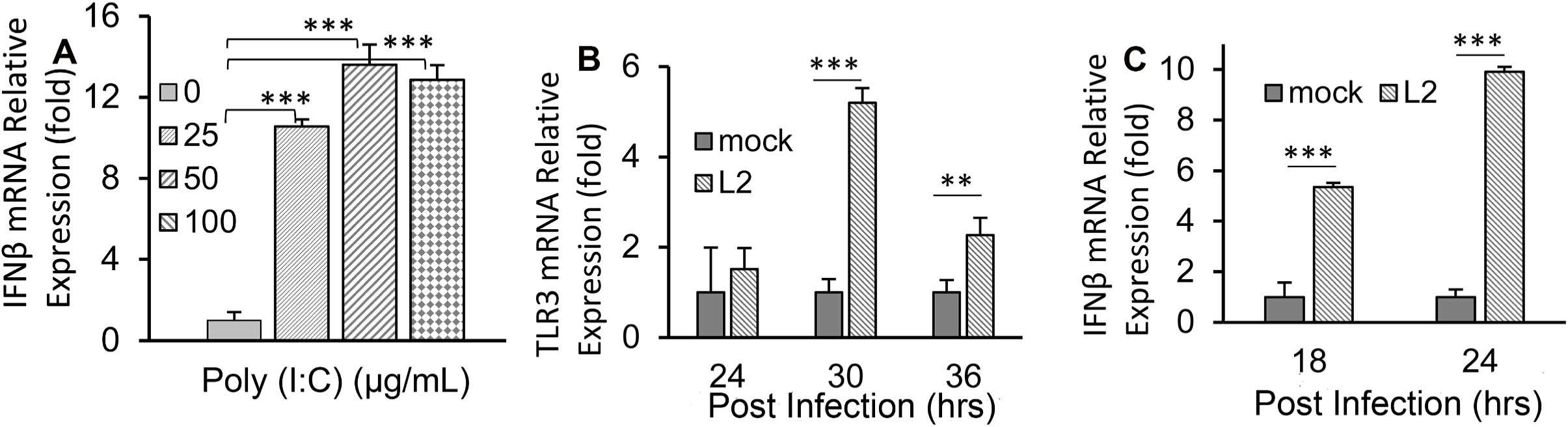
TLR3 is present and functional in human OE-E6/7 cells. Human OE-E6/7 cells were seeded in 12 well plates and cultured in DMEM only, or DMEM supplemented with either poly (I:C) for 24hrs or infected with 10 IFU/ cell *C. trachomatis*-L2 for 36hrs. (A) Poly (I:C) induced the expression of IFN-β mRNA in a dose-dependent manner. Relative expression of (B) IFN-β mRNA and (C) TLR3 mRNA in response to *C. trachomatis*-L2 infection at the time-point indicated. Data are representative of at least three independent experiments.

### Disruption of TLR3 function in human genital tract epithelial cells and clone identification

To ascertain the role of TLR3 in the innate-immune response to human genital tract *Chlamydia* infections *in vitro*, we disrupted TLR3 function in both human oviduct (hOE cells) and cervical (Hela cells) tissue using the Sigma CRISPR Lentivirus system (see Materials and Methods). The CRISPR system consisted of 3 gRNA sequences which targeted human TLR3 gene at exon 2, exon 2, and exon 4 respectively (Fig. S1). To ascertain the efficacy of using this approach to disrupt TLR3 function in these cells, we first examined the TLR3 protein expression levels in the putative clones by using capillary electrophoresis in the Wes™ system to identify and quantitate TLR3 protein. In these experiments, we loaded a total 400ng of cell lysate isolated from each of the selected clones and the WT control cells. Figure 2 shows the results of capillary electrophoresis in which we simultaneously immunoassayed for both TLR3 expression and the β-Actin loading control in hOE cells, Hela cells, and the representative TLR3-deficient clones generated in each parental cell-line. As shown, a major band indicative of the TLR3 protein expression was identified in the hOE and Hela cells, with a peak molecular weight of 174kDa and 187kDa, respectively. Our data are suggestive that the TLR3 protein is post-translationally modified in different ways in the different cell types, which affects their electrophoretic migration and apparent molecular weight (37).

**FIGURE 2.**
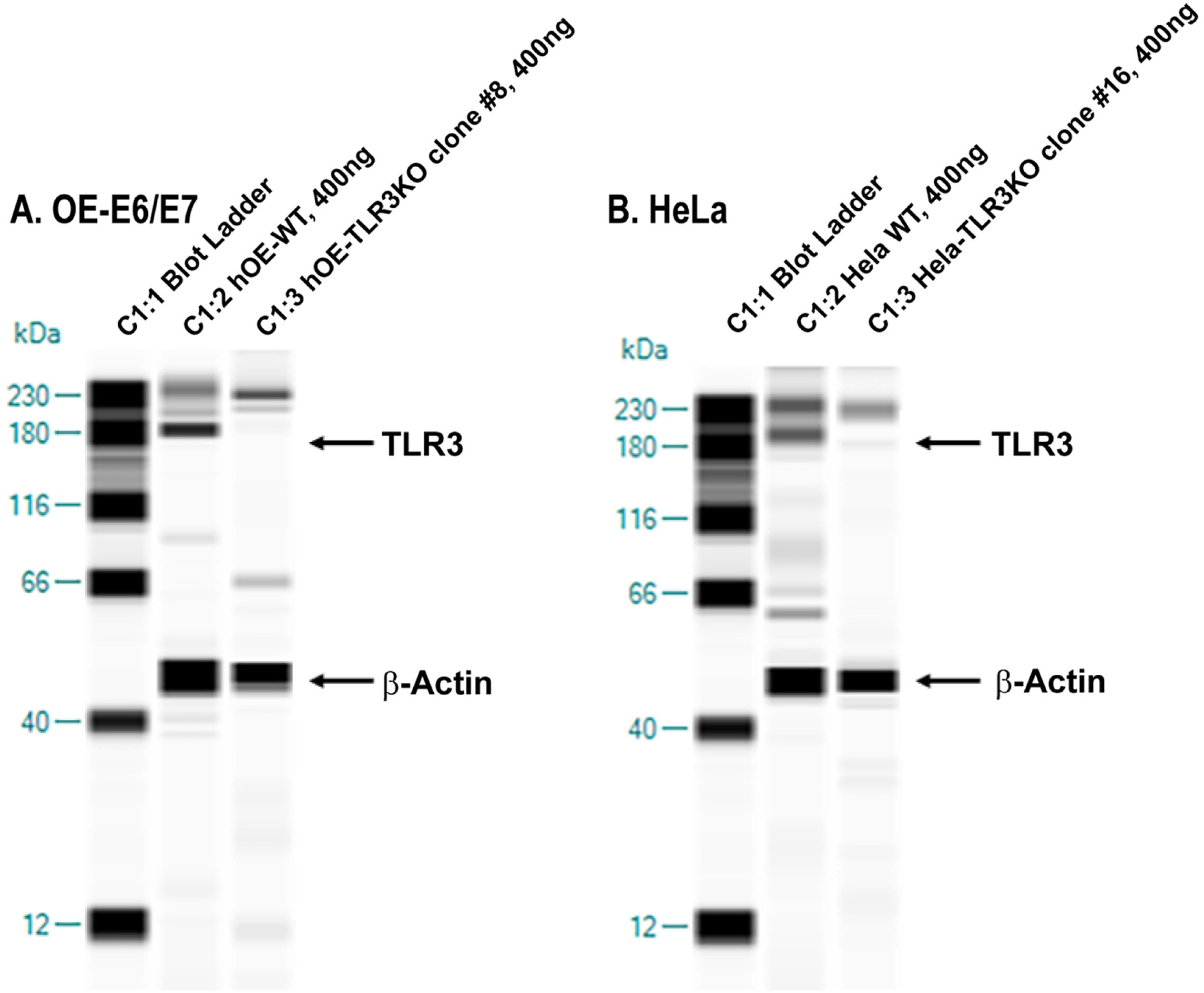
Disruption of TLR3 protein expression in human oviduct and cervical cells. Wes™ simple western system was used to confirm the disruption of TLR3 protein expression: (A) <u>Human OE-E6/7 cells.</u> Lane 1, Ladder; lane 2, 400ng WT human OE-E6/7 cell protein; lane 3, 400ng hOE-TLR3KO cell protein. (B) <u>Hela cells.</u> Lane 1, Ladder; lane 2, 400ng WT Hela cell protein; lane 3, 400ng Hela-TLR3KO cell protein. Data presented are representative data of selected TLR3-deficient clones from both cell types.

A qualitative examination of Figure 2 shows that TLR3 protein expression levels were substantially lower in both clone #8 of the hOE-TLR3KO cells and clone #16 of the Hela TLR3KO cells when compared to their wild-type counterparts. These clones were selected and used as TLR3-deficient versions of the human OE-E6/7 cells and the Hela cells, respectively, and were quantitatively analyzed for actual TLR3 expression levels in the Wes™ Compass software. To ascertain the actual level of reduction in TLR3 protein expression levels, the protein band peaks were identified and quantified using the chemiluminescence peak area of the Wes™ generated data. The ratio of TLR3 protein expression compared to that of the β-Actin loading control was used as the index for calculating relative TLR3 protein expression level. In the hOE cells, the ratios of TLR3/β-Actin for WT and TLR3KO were 5732/41538=0.138 and 347/16533=0.021, respectively. When calculated, the TLR3 protein expression was down about 85% in clone #8 of the hOE-TLR3KO cells. In Hela cells, the ratio of TLR3/β-Actin for WT and TLR3KO was 9254/43601=0.212 and 310/16482=0.018, respectively. The calculated expression level of TLR3 protein was down about 94% in clone #16 of the Hela TLR3KO cells. Collectively, our findings demonstrate that CRISPR Lentivirus system was very effective in disrupting TLR3 protein expression in both of these cell types.

### TLR3 deficiency in human genital tract epithelial cells results in a decreased IFN-β synthesis in response to poly (I:C) stimulation and chlamydial infection

To determine whether the TLR3-deficient clones representing human oviduct and cervical tissue were also deficient in their ability to elicit appropriate TLR3-dependent innate-immune responses, we first treated the clone representative of each cell type with 0, 25, 50, 100 µg/mL the TLR3 agonist poly (I:C) for 24hrs and assessed TLR3’s functionality by measuring the induction of IFN-β gene expression (Figure 3). As shown in Figure 3A, the hOE TLR3KO cells exhibited significantly lower levels of IFN-β gene expression in response to poly (I:C) induction at all concentrations tested when compared to the non-target CRISPR control cells. We observed similar reductions in the induction of IFN-β transcription in response to poly (I:C) in the representative Hela TLR3KO clones (Figure 3B). However, the fold-difference in IFN-β gene induction between the TLR3-deficient clones and the non-target CRISPR control for the Hela cells seemed to decrease at the higher poly (I:C) concentrations. We observed no noticeable differences in poly (I:C) induction of IFN-β transcription between the WT cells and their non-target CRISPR control counterpart for either cell type *(data not shown).* These findings show a successful disruption of TLR3 function in the TLR3-deficient epithelial cell clones representing human oviduct and cervical tissues.

**FIGURE 3.**
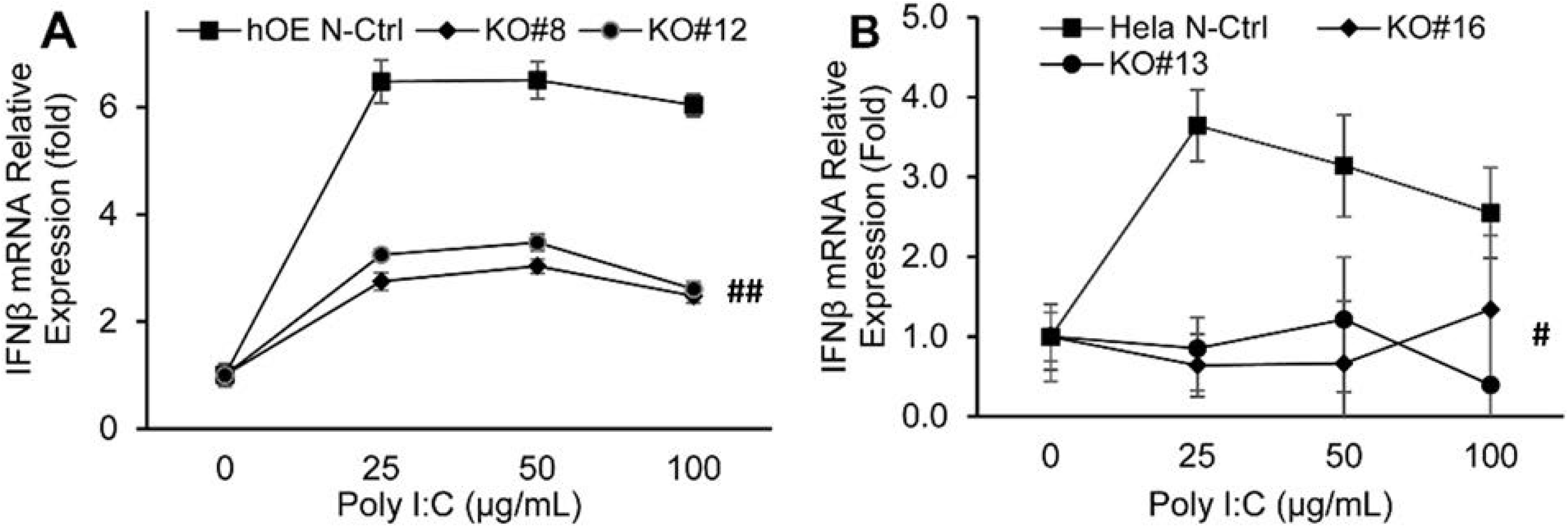
Disruption of TLR3 function by CRISPR dramatically decreases IFN-β mRNA expression in response to poly (I:C) stimulation. Selected clones representing TLR3-deficient: (A) Oviduct [hOE] and (B) Cervical [Hela] cells were treated with 0, 25, 50, 100 µg/mL poly (I:C) for 24hrs. The cells were harvested for total cell RNA isolation. The response to poly (I:C) was determined by measuring the induction of IFN-β mRNA synthesis via qPCR. The relative gene expression levels of each clone compared to their respective non-target CRISPR controls are shown. Data are representative of at least three independent experiments.

In order to make comparisons to what we have previously reported on the role of TLR3 in the innate immune response to *Chlamydia* infection in the murine oviduct epithelial cells, we put more emphasis on the hOE cells in this study and selected the hOE TLR3KO clone #8 for use in the remainder of this report. We first sought to ascertain the impact of TLR3 deficiency on the *C. trachomatis*-induced synthesis of IFN-β in the human oviduct epithelial cells. We infected hOE-WT, hOE N-Ctrl, and hOE-TLR3KO cells with 10 IFU *C. trachomatis*-L2 before harvesting cell supernatants at 18 and 36h post-infection to measure the amount of IFN-β secreted. Figure 4A shows a significant reduction in the *Chlamydia*-induced synthesis of IFN-β in the TLR3-deficient hOE cells when compared to both the WT and non-template control cells. Our data show a 40-50% reduction in the amount of IFN-β synthesized at both time-points and are suggestive that TLR3 plays a significant role in the optimal synthesis of IFN-β in *C. trachomatis* infected hOE cells.

**FIGURE 4.**
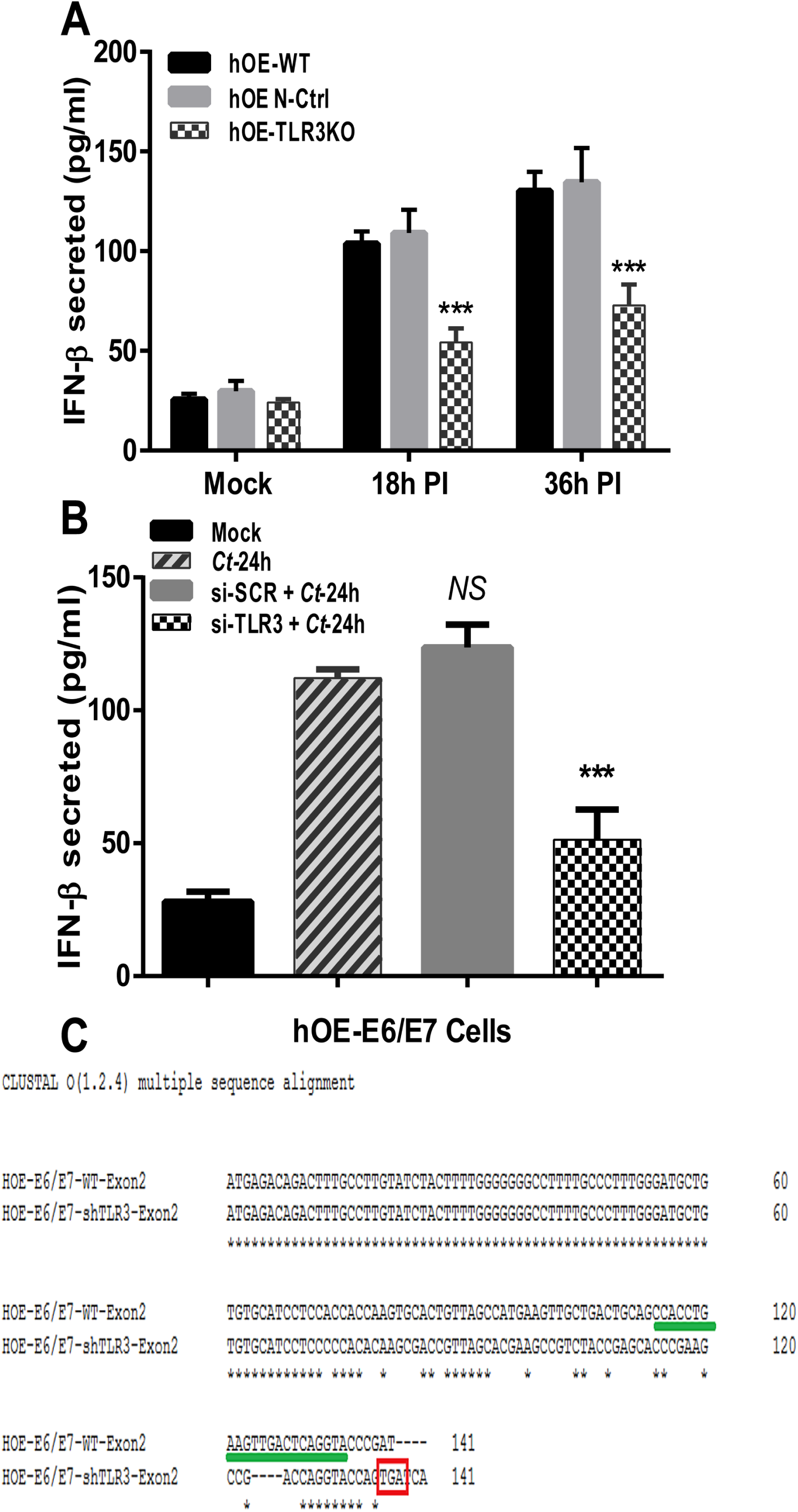
TLR3 deficiency results in the decreased synthesis of IFN-β during *C. trachomatis* infection of oviduct epithelium. (A) WT and TLR3-deficient hOE cells were either mock infected or infected with *C. trachomatis*-L2 at a MOI 10 IFU/ cell for up to 36hrs. Supernatants were collected from cells representing mock, 18h, and 36h post-infection before measuring the chlamydial induced synthesis of IFN-β by standard ELISA. (B) WT hOE cells were infected with 10 IFU/cell *C. trachomatis*-L2 24hrs after treatment with 2.5 µg/ml of either si-SCR or TLR3-specific siRNA (si-TLR3). Supernatants were harvested at 24hrs post-infection to measure the chlamydial induced synthesis of IFN-β by standard ELISA. (C) CLUSTAL-O multiple sequence alignment of a sequenced PCR product containing exon #2 of human TLR3 reveal early stop codon (TGA) at position 141. The TGA stop codon is inside the red box, and gRNA sequences are highlighted with a green line. Data are representative of at least 3 independent experiments. Statistically significant differences are shown by asterisks (***, *p*<0.005).

We next assessed whether our limiting dilution clonal expansion procedure introduced ‘founder effects’ such as causing a global dysregulation in TLR signaling that leads to the diminished synthesis of inflammatory cytokines during *Chlamydia* infection in hOE TLR3KO cells. To test whether there were founder effects introduced that negatively impacted TLR signaling in a global manner, we treated WT and TLR3-deficient hOE cells with ultrapure preparations of ligands for TLR2 (peptidoglycan), TLR5 (flagellin), and TLR9 (ODN-CpG) for 24hrs before testing supernatants for IL-6 synthesis by ELISA (14). As shown in Fig. S2, we saw no significant differences between WT and TLR3-deficient hOE cells in the synthesis of IL-6 when the cells were treated with agonists for either TLR2, TLR5, or TLR9. To address the possibility that reduced IFN-β synthesis observed in the hOE-TLR3KO cells was due to pathways unrelated to TLR3 that may have been disrupted by CRISPR, we used siRNA as an alternative method to transiently disrupt TLR3 gene expression and protein function in WT hOE cells. Figure 4B demonstrates significant reductions in the amount of IFN-β secreted into the supernatants of *C. trachomatis*-L2 infected hOE cells that were pretreated with TLR3-specific siRNA 24hrs prior to infection.

To discern the exact nature of the TLR3 gene defect introduced by CRISPR-Cas9, clone #8 of the hOE-TLR3KO cells was selected for gene sequencing and gene sequence alignment in CLUSTAL-O. Analysis of the TLR3 gene sequencing data revealed that clone #8 of the hOE-TLR3KO cells was heterozygous for TLR3 gene disruption at exon #2 and that the resultant knockout allele incorporated a premature stop codon (TGA) at nucleotide position 141 (Figure 4C). The CRISPR-Cas9 method of gene disruption resulted in the translation of truncated TLR3 protein that is likely non-functional, and our collective data examining TLR3 protein expression and function (Figs 2-4 and Fig. S2) are suggestive that there is a minimal contribution from the TLR3 allele that was not disrupted by CRISPR-Cas9 at Exon #2. Collectively, these findings show that CRISPR-Cas9 methodology was effective in disrupting TLR3 gene function in hOE cells, and corroborate our previous reports that implicate TLR3 as a major contributor to the *Chlamydia*-induced synthesis of IFN-β during *Chlamydia* infection of oviduct epithelial cells (13, 38).

### TLR3 deficiency alters the *Chlamydia*-induced syntheses of acute inflammatory biomarkers in human oviduct epithelial cells

We previously reported that murine OE cells showed dysregulation in the *C. muridarum* induced syntheses of a multitude of acute inflammatory mediators, and subsequently showed that TLR3 deficient mice exhibited increased chlamydial shedding, aberrant T-cell recruitment, and more severe genital tract pathology when compared to WT mice (15, 32). To assess whether the absence of TLR3 is associated with the chlamydial-induced syntheses of key biomarkers of inflammation in human oviduct epithelial cells, we performed multiplex ELISA analysis for the detection and quantification of key human innate-immune inflammatory biomarkers. hOE-WT and hOE-TLR3KO cells were either mock-infected, infected with *C. trachomatis*-L2 for up to 30hrs, or infected with *C. trachomatis*-serovar D for up to 72hrs before multiplex analyses of the cell supernatants. As shown in Figures 5 and 6, TLR3-deficient hOE cells secreted significantly reduced levels of the acute inflammatory markers IL-6, IL-6Ra, sIL-6Rβ (gp130), IL-8, IL-20, IL-26, and IL-34 secreted into the supernatants of the *C. trachomatis* infected cells throughout infection. Conversely, TLR3 deficiency in hOE cells resulted in the *increased* synthesis of CCL5 and the type III interferon IL-29 (IFNλ1) throughout infection, and IL-28A (IFNλ2) late in infection (Figures 6 and 7). Collectively, these data demonstrate a putative role for TLR3 in the acute phase of the innate immune response to *Chlamydia* that is known to occur early during infection *in vivo* (*17, 39–42*), and that TLR3 has some role in modulating mediators of the adaptive immune response (43).

**FIGURE 5.**
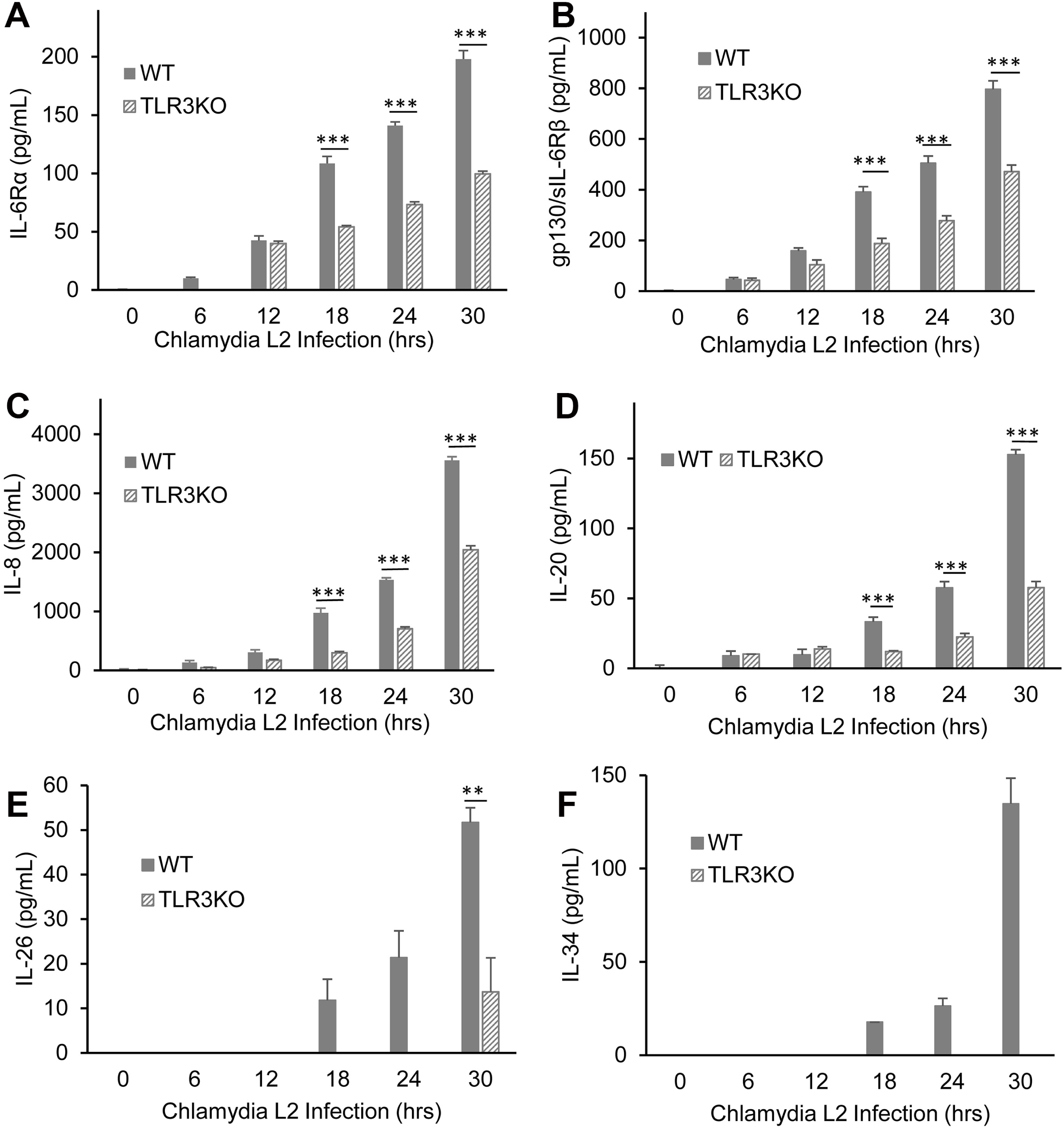
TLR3 deficiency results in the attenuation of many of the acute-phase inflammatory mediators. WT and TLR3-deficient hOE cells were infected with *C. trachomatis*-L2 at a MOI 10 IFU/ cell for up to 30hrs. Supernatants were collected from individual wells every 6 hours for multiplex ELISA analyses to measure the expression of: (A) IL-6Ra; (B) sIL-6Rβ (gp130), (C) IL-8, (D) IL-20, (E) IL-26, and (F) IL-34. Statistically significant differences are shown by asterisks (**, *p*<0.01; ***, *p*<0.001). Data are representative of three independent experiments.

**FIGURE 6.**
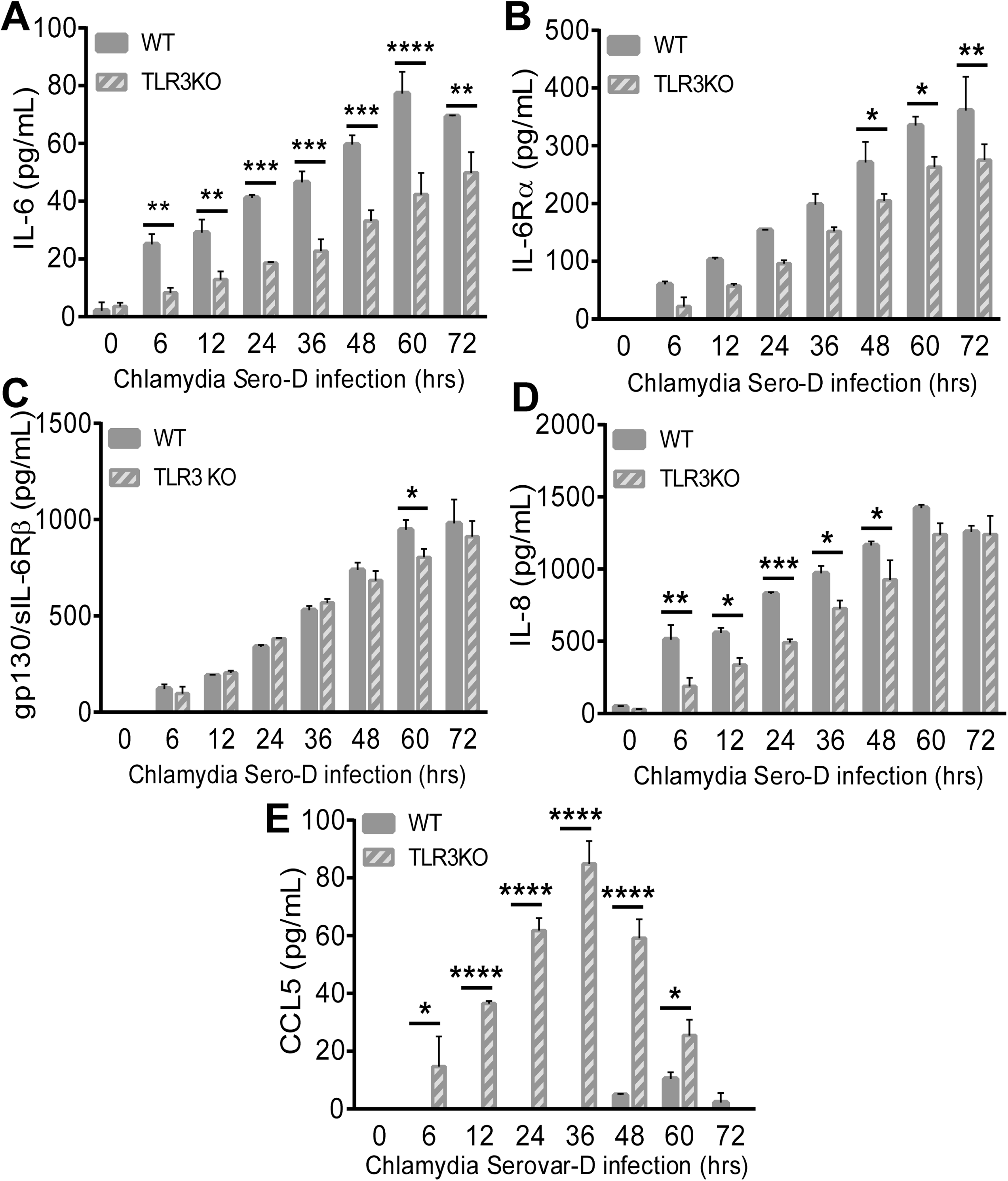
TLR3 deficiency alters the acute-phase inflammatory mediator synthesis during infection with *C. trachomatis*-serovar D. WT and TLR3-deficient hOE cells were infected with *C. trachomatis*-serovar D at a MOI 10 IFU/ cell for up to 72hrs. Supernatants were collected from individual wells at the time listed for multiplex ELISA analyses to measure the expression of: (A) IL-6, (B) IL-6Rα, (C) sIL-6Rβ (gp130), (D) IL-8, and (E) CCL5. Statistically significant differences are shown by asterisks (*, *p*<0.05; **, *p*<0.01; ***, *p*<0.005). Data are representative of three independent experiments.

**FIGURE 7.**
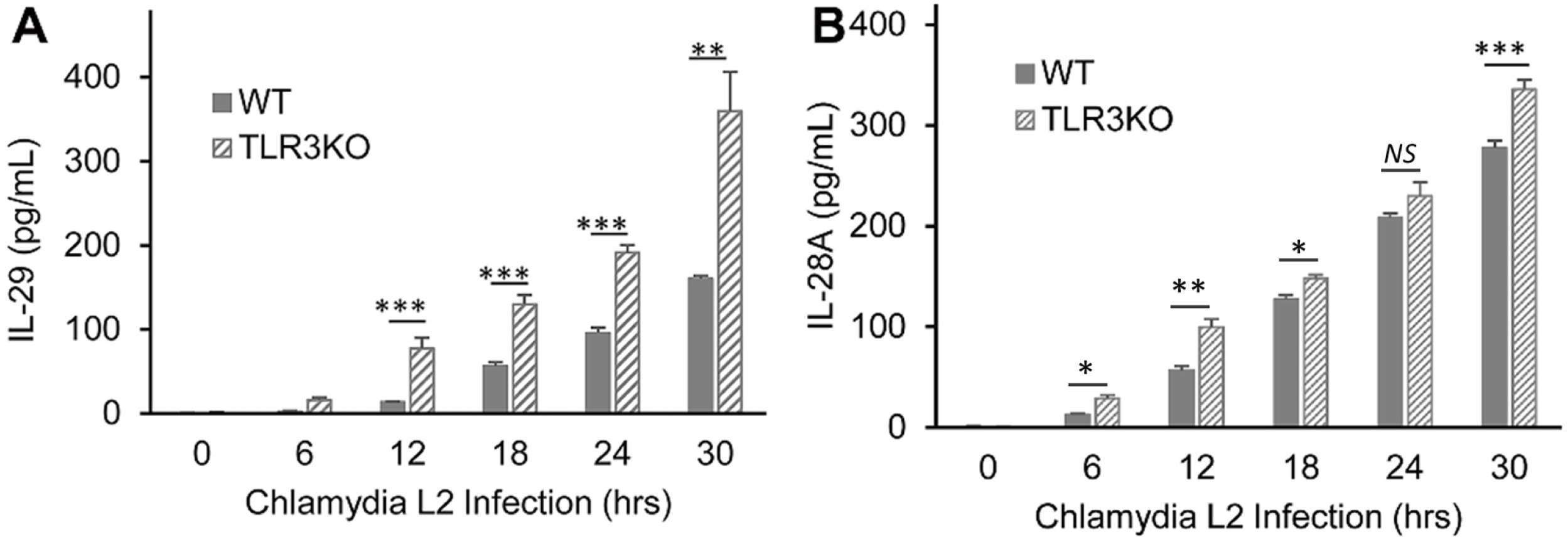
TLR3 deficiency causes the increased expression of type-III IFNs. WT and TLR3-deficient hOE cells were infected with *C. trachomatis*-L2 at a MOI 10 IFU/ cell for up to 30hrs. Supernatants were collected from individual wells every 6 hours for multiplex ELISA analyses to measure the expression of: (A) IL-29 and (B) IL-28A. Statistically significant differences are shown by asterisks (**, *p*<0.01; ***, *p*<0.001). Data are representative of three independent experiments.

### TLR3 has a functional role in the *C. trachomatis* induced syntheses of factors associated with genital tract fibrosis, scarring, and chronic inflammation

In our previous investigations into the role of TLR3 in the pathogenesis of genital infections in mice, one key aspect of TLR3 deficiency that we observed was that mice deficient in TLR3 appeared exhibit indicators of more pronounced chronic sequelae, such as lymphocytic endometritis and hydrosalpinx (32). To examine whether TLR3 has a similar role in the pathogenesis of genital tract *Chlamydia* infections in humans, we next measured the *Chlamydia*-induced synthesis of biomarkers associated with chronic inflammatory disease and tissue necrosis in the WT and TLR3 deficient hOE cells. As shown in Figures 8A and 8D, *C. trachomatis* infection induces the synthesis of soluble tumor necrosis factor receptor 1 (sTNF-R1) in both cell types; however, its synthesis was significantly lower in the TLR3-deficient hOE cells at mid-to-late times during infection. The exact role of sTNF-R1 in *Chlamydia* infection is not clear, but it is a known negative regulator of TNFα, which is a cytokine associated with severe genital tract sequelae in mice (44). Figure 8B shows that TLR3 deficiency leads to a similar impact on the chlamydial induced synthesis of tumor necrosis factor ligand superfamily member 13B (TNFSF13B), a cytokine that belongs to the tumor necrosis factor family that acts as a potent B-cell activator (45). Interestingly, Figures 8C and 8E shows that the syntheses of TGF-β1 and ICAM-1 is increased during *C. trachomatis* infection of hOE-TLR3KO cells, implicating TLR3 in the negative regulation of key components of the pathophysiological process of fibrogenesis (46).

**FIGURE 8.**
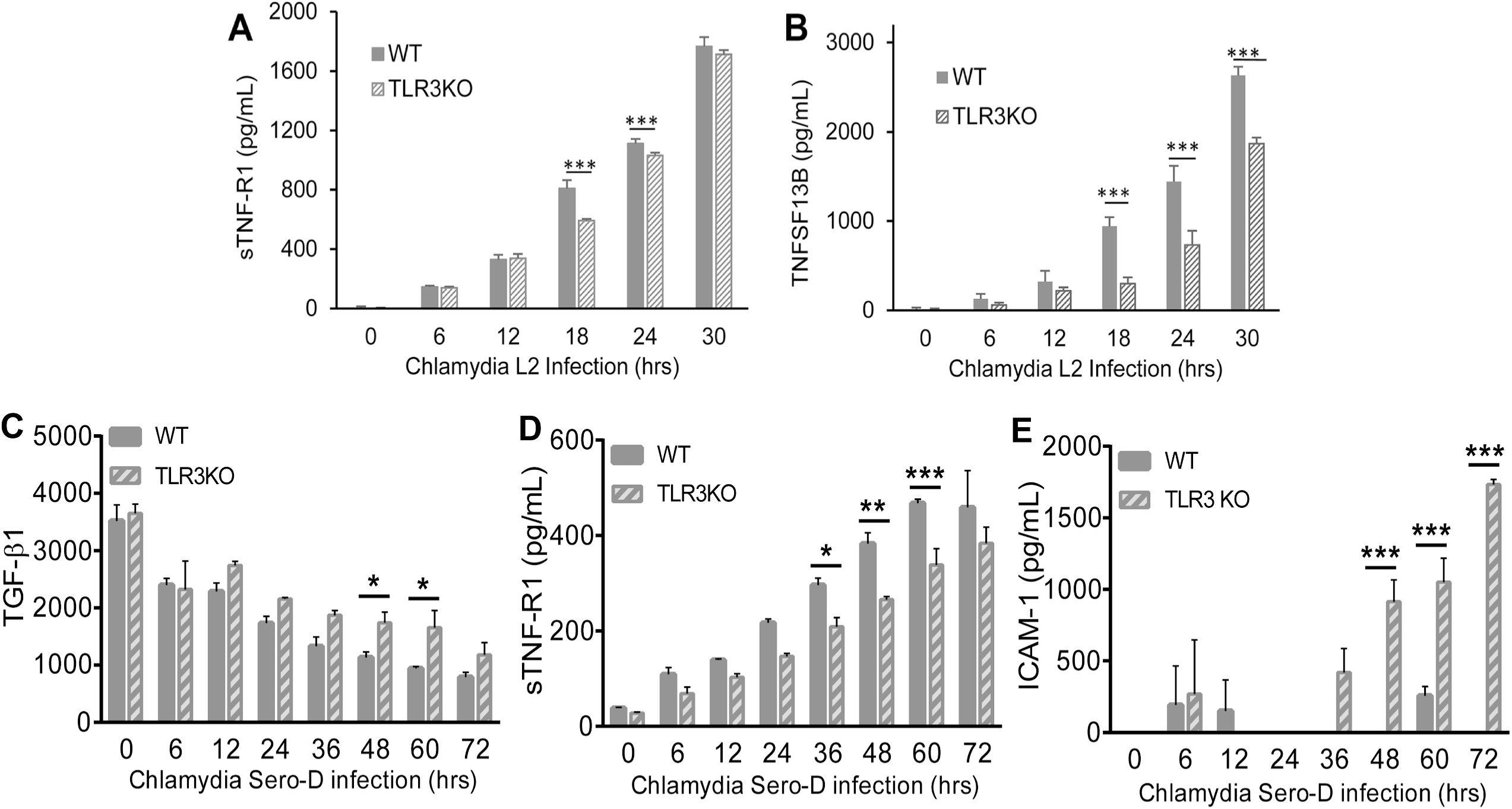
TLR3 deficiency alters the synthesis of cytokines associated with cellular adhesion and tissue integrity during genital tract infection with *C. trachomatis*. WT and TLR3-deficient hOE cells were infected with either *C. trachomatis*-L2 (**A-B**) or *C. trachomatis*-serovar D (**C-E**) at a MOI 10 IFU/ cell for up to 72hrs. Supernatants were collected from individual wells at the time listed for multiplex ELISA analyses to measure the expression of: (**A, D**) sTNF-R1, (**B**) (TNFSF13B), (**C**) TGF-β1, and (**E**) ICAM-1. Statistically significant differences are shown by asterisks (*, *p*<0.05; **, *p*<0.01; ***, *p*<0.005). Data are representative of three independent experiments.

The matrix metalloproteinases (MMPs) are a tightly regulated family of proteins that are involved in the breakdown of extracellular matrix in normal physiological processes and are known to have a regulatory role in the inflammatory immune response, wound healing, cell migration, and embryonic development (47). Dysregulation of these proteins during *Chlamydia* infection has been demonstrated to play a role in the pathogenesis of fallopian tube damage during genital tract infections, and in corneal scarring in patients with trachoma (48, 49). To ascertain whether MMPs are synthesized in response to *C. trachomatis* infection in the hOE cells, and to determine whether TLR3 deficiency impacts their protein expression levels, we next measured the secretion of candidate MMPs into the supernatants of *C. trachomatis* infected cells in our multiplex ELISA. As shown in Figure 9, *Chlamydia* infection induced the production of MMP-1, MMP-2, MMP-3, and MMP-10 throughout infection, supporting the observations of others regarding the induction of MMPs in genital tract infections (48, 50–52). The *Chlamydia*-induced syntheses of these MMPs were either completely absent or severely diminished in their production in the TLR3-deficient hOE cells relative to hOE-WT cells when infected with the L2 serovar (Figures 8A-8C). However, the synthesis levels were more similar when infected with serovar D, albeit at significantly lower amounts in the hOE-TLR3KO cells at various points post-infection (Figures 8D-8F). Taken together, data from Figures 8 and 9 are suggestive that TLR3 plays some role in regulating the syntheses of immune factors involved in modulating the genital tract pathologies associated with *Chlamydia* infection in humans.

**FIGURE 9.**
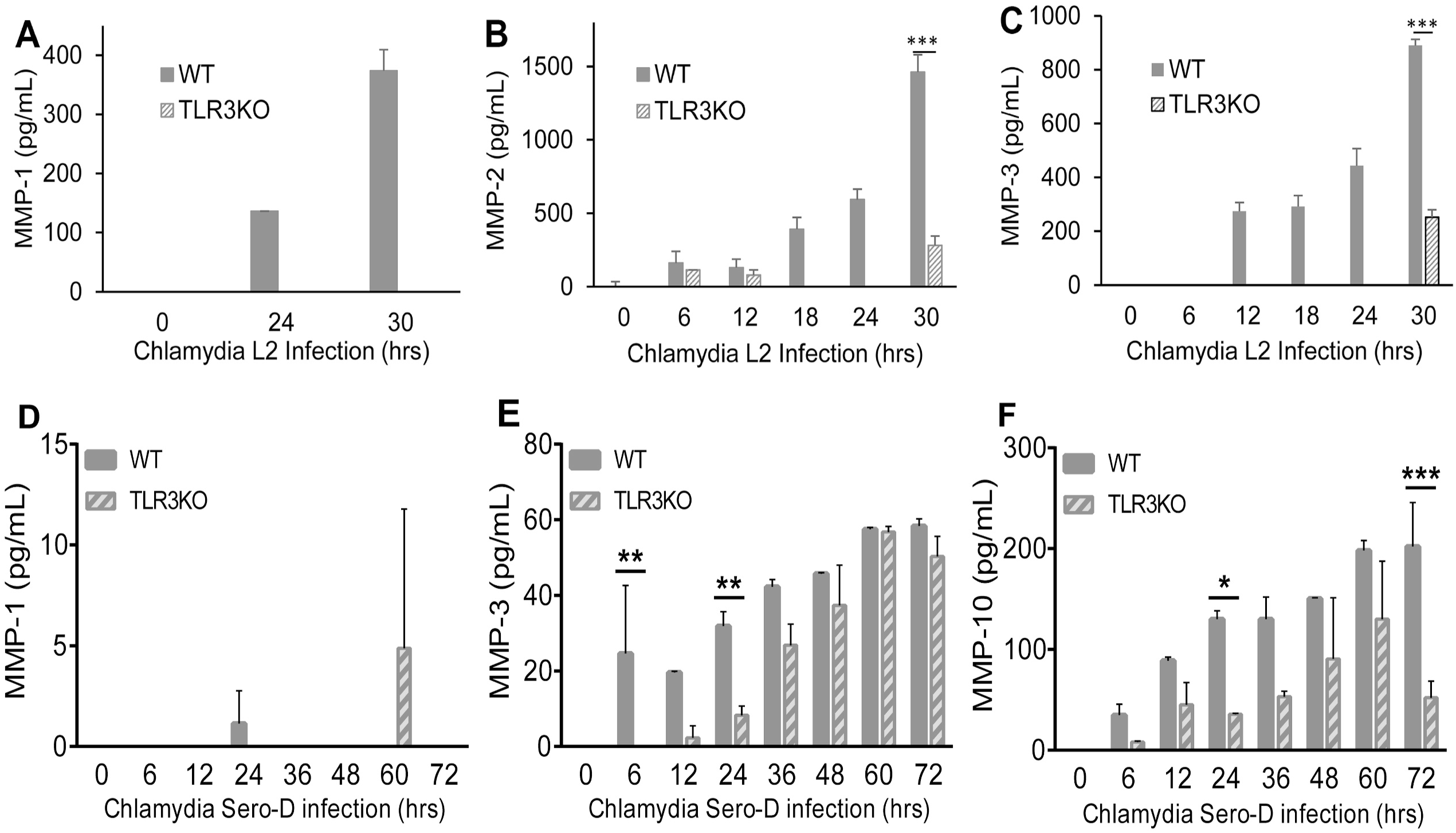
TLR3 regulates the *Chlamydia*-induced expression of matrix metalloproteinases during genital tract infections with *C. trachomatis*. WT and TLR3-deficient hOE cells were infected with 10 IFU/ cell with either *C. trachomatis*-L2 (**A-C**) or *C. trachomatis*-serovar D (**D-F**) for up to 72hrs. Supernatants were collected from individual wells at the time listed for multiplex ELISA analyses to measure the expression of: (**A, D**) MMP-1, (**B**) MMP-2 (**C, E**) MMP-3, and (**F**) MMP-10. Statistically significant differences are shown by asterisks (*, *p*<0.05; **, *p*<0.01; ***, *p*<0.005). Data are representative of three independent experiments.

### TLR3 signaling regulates the *Chlamydia*-induced synthesis of biomarkers associated with persistence, metastasis, and autoimmunity

A major factor in the protective immune response to *C. trachomatis* infection is the synthesis of gamma-interferon (IFNγ), which inhibits the growth of chlamydial RBs via mechanisms that lead to tryptophan starvation, chlamydial death, and eventual clearance of the bacteria (53). However, recent evidence of chlamydial reticulate bodies being able to substantially alter their gene transcription, decrease metabolism, and entering into what is known as a ‘persistent’ state, suggests a survival mechanism that *Chlamydia* has evolved to evade immune surveillance (54–56). Persistence in microbial infections are often implicated in the triggering of autoimmune reactions, and this was demonstrated in studies investigating the role that *Chlamydia* persistence plays in triggering self-immune reactions in infected male rodents (57).

To determine whether *Chlamydia* infection induces a cellular response that signals a state of persistence in the infected hOE cells, our multiplex analyses included several biomarkers for persistence and autoimmunity that are known to be secreted by cells in various clinical syndromes. Figures 10 and 11 show the results of the chlamydial induction of soluble CD163, chitinase-3-like 1, Lipocalin-2, osteopontin, and pentraxin-3 throughout infection in WT and TLR3 deficient hOE cells. As shown, *Chlamydia* induces the synthesis of sCD163 (a factor associated with long-term chronic inflammatory diseases such as rheumatoid arthritis) and the anti-apoptotic chitinase-3-like 1 protein in the hOE cells. However, our data show that the protein secretion levels of these biomarkers are significantly reduced in the absence of TLR3 when these cells were infected with the L2 serovar. Although secretion levels of sCD163 were much higher during infection with serovar D, there was no significant reduction during TLR3 deficiency as was observed in the L2 infection.

Figures 10 and 11 show that TLR3 deficiency leads to significantly *increased* levels of osteopontin and pentraxin-3. The role of these proteins in *Chlamydia* infection is not clear; however, osteopontin is an inflammatory mediator often associated with autoimmune disease, chronic inflammatory disorders, and progression of tumor cells (58), while pentraxin-3 is a known biomarker for pelvic inflammatory disease (PID) and is also associated with autoimmune diseases (59, 60). Collectively, our findings indicate that TLR3 may have a functional role in regulating the expression of biomarkers symbolic of long-term or persistent disease states in the *Chlamydia*-infected hOE cells, which is a precondition that often correlates with the initiation of autoimmune responses and chronic sequelae (61–64).

**FIGURE 10.**
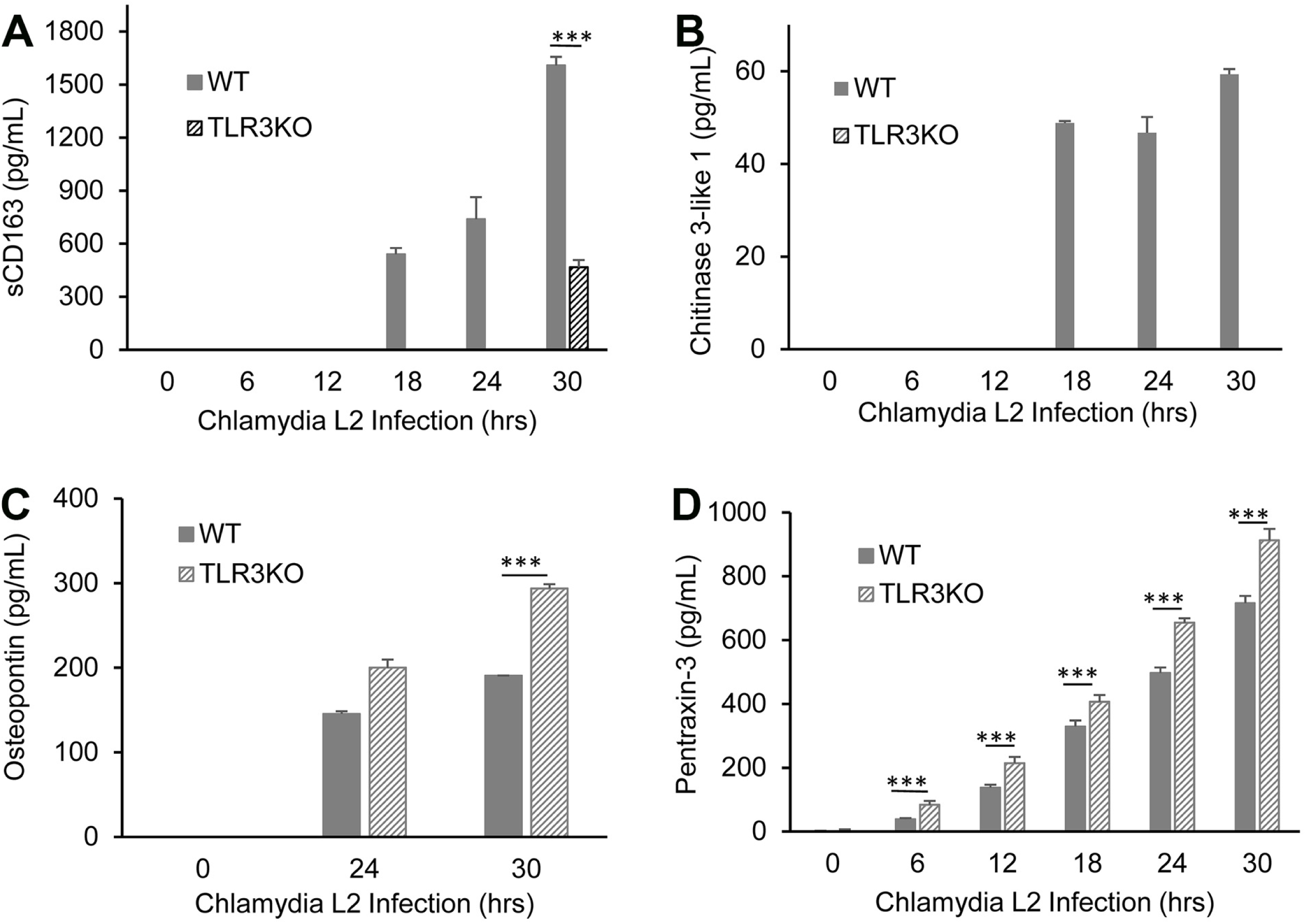
TLR3 plays a role in regulating the *Chlamydia*-induced syntheses of biomarkers associated with persistence and autoimmunity. Secreted protein levels of (A) sCD163, (B) Chitinase-3-like 1, (C) Osteopontin, and (D) Pentraxin-3 were measured in the supernatants of WT and TLR3-deficient hOE cells that were infected with *C. trachomatis*-L2 at a MOI 10 IFU/ cell for up to 30hrs. The supernatants were collected from individual wells for multiplex ELISA analyses at the listed time point post-infection. Statistically significant differences are denoted by asterisks (***, *p*<0.001). Data are representative of three independent experiments.

**FIGURE 11.**
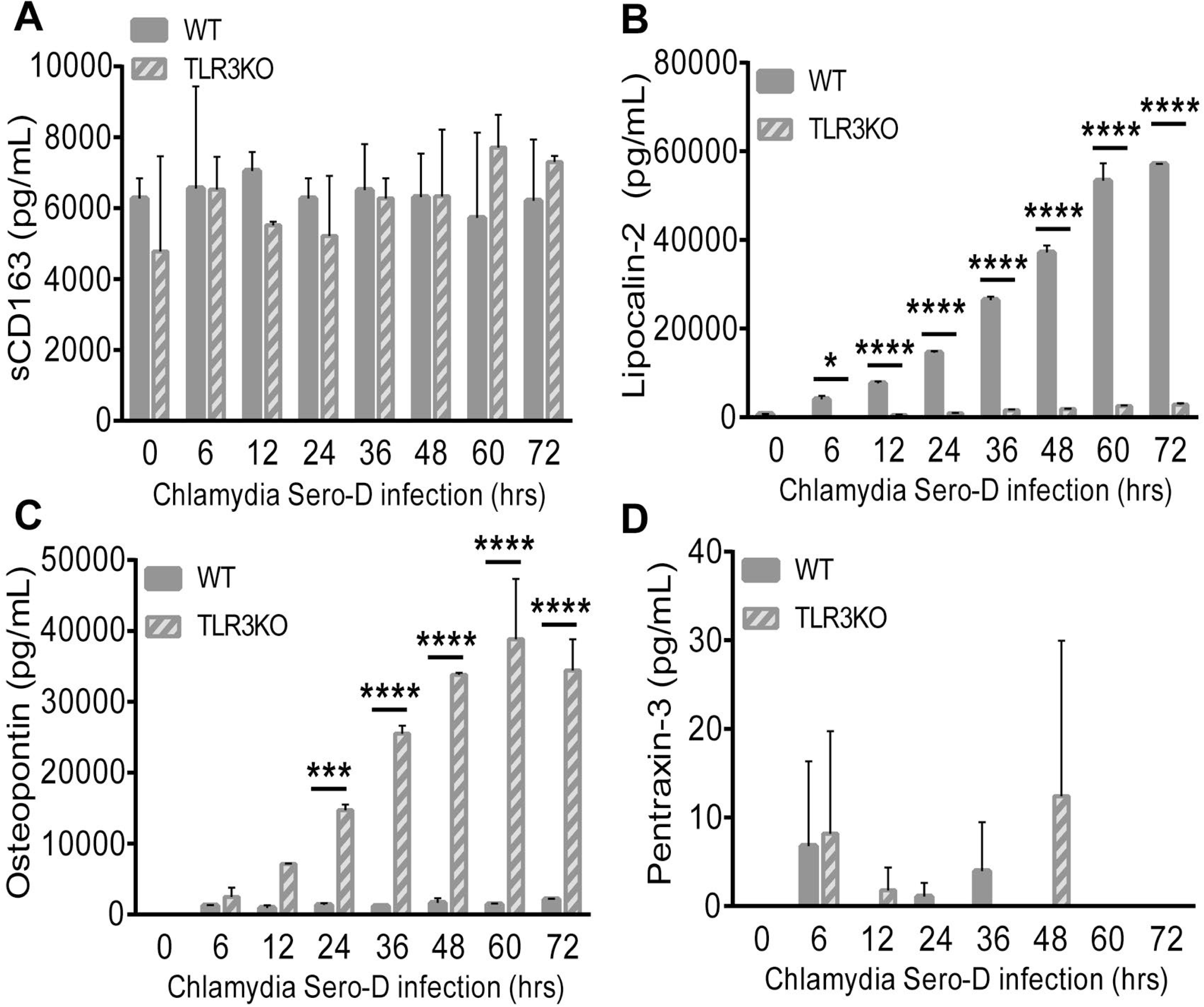
TLR3 plays a role in regulating the *Chlamydia*-induced syntheses of biomarkers associated with iron sequestration, persistence, and autoimmunity during infection with *C. trachomatis*-serovar D. WT and TLR3-deficient hOE cells were infected with *C. trachomatis*-serovar D at a MOI 10 IFU/ cell for up to 72hrs. Supernatants were collected from individual wells at the time listed for multiplex ELISA analyses to measure the expression of: (A) sCD163, (B) Lipocalin-2, (C) Osteopontin, and (D) Pentraxin-3. Statistically significant differences are shown by asterisks (*, *p*<0.05; ***, *p*<0.005; ****, *p*<0.001). Data are representative of three independent experiments.

### TLR3 deficiency altered the LPS content and size of the chlamydial inclusion

Our previous investigations into mechanisms that impact the chlamydial-induced synthesis of IFN-β showed that disruption of IFN-β had a significant impact on the chlamydial inclusion size and chlamydial replication. In that regard, we showed that *C. muridarum* replication in murine OE cells deficient in either TLR3 or STAT-1 was more robust and that the inclusions were larger and aberrantly shaped [(15, 65); Fig S3]. To determine the physiological consequences of TLR3 deficiency on *Chlamydia* inclusion size in human oviduct epithelial tissue, we infected WT and TLR3-deficient hOE cells with *C. trachomatis*-serovar D at a MOI of 10 IFU/ cell for 36hrs. We next stained the cells for chlamydial LPS using the EVI-HI anti-*Chlamydia* LPS antibody, and examined the infected cells for chlamydial inclusion size and LPS content in fluorescent microscopy and multi-spectral flow cytometry, respectively. We first examined *C. trachomatis*-serovar D infected hOE-WT and hOE-TLR3KO cells by immunofluorescent microscopy to get a qualitative comparison of the chlamydial inclusion size in order to ascertain whether TLR3 has any impact on *Chlamydia* development. As demonstrated in the representative capture shown in Figure 12, we routinely saw that the chlamydial inclusions in the TLR3-deficient hOE cells were much larger, amorphously shaped, and were more diffusely stained with punctate patterns throughout the inclusion. In contrast, the inclusions in the hOE-WT cells were generally more compact in size, more uniformly stained, and had more staining intensity per pixel than the TLR3-deficient hOE cells. We saw very similar trends in the *C. muridarum*-infected WT and TLR3(-) OE cells derived from mice (Fig. S3).

**Figure 12.**
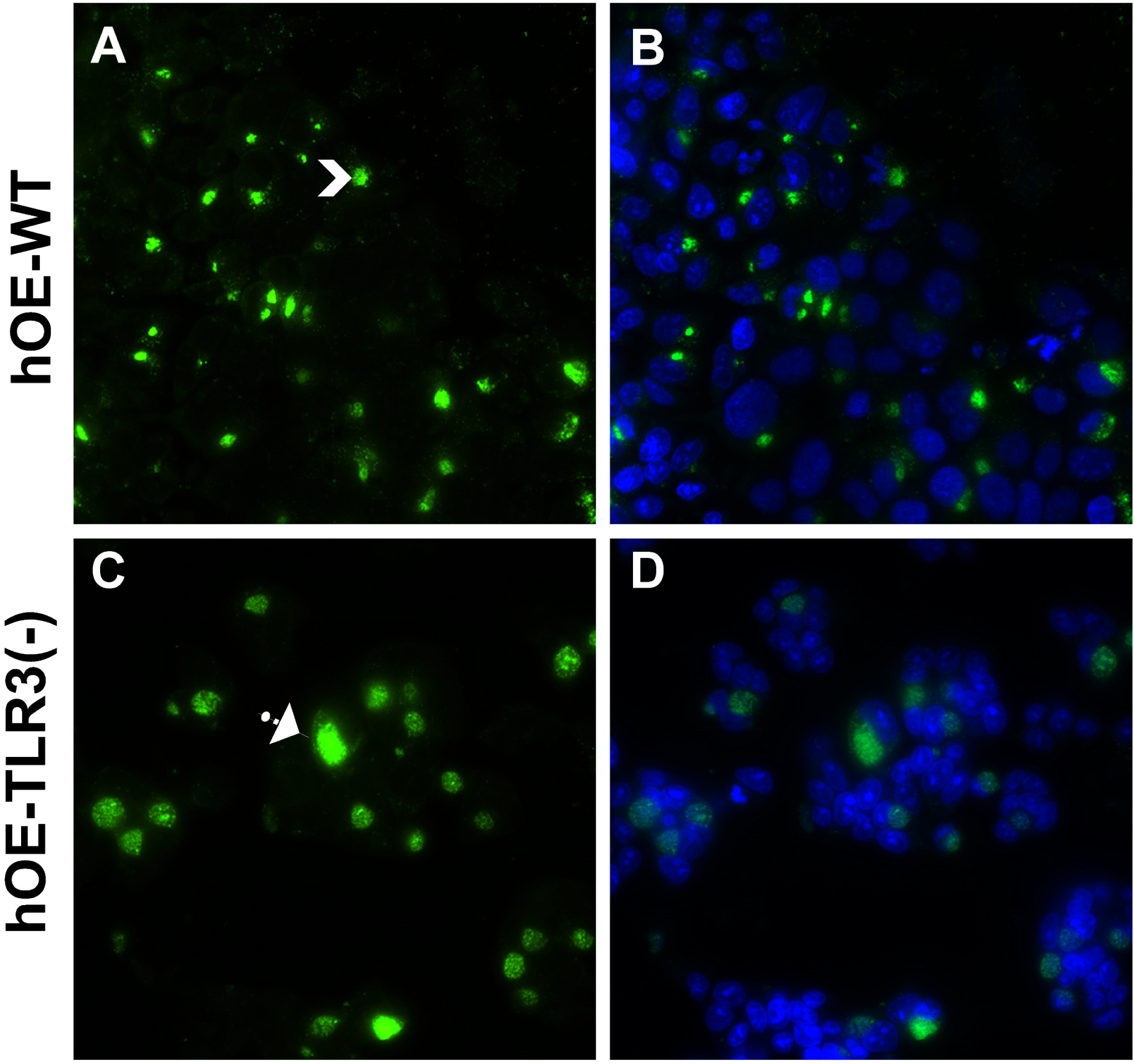
TLR3 deficiency in murine OE cells leads to larger and aberrantly shaped chlamydial inclusions. hOE-WT cells (A and B) and hOE-TLR3KO cells (C and D) were either mock treated or infected with *C. trachomatis*-serovar D at a MOI of 10 IFU/ cell for 36hrs. The chlamydial inclusion was stained using anti-chlamydial LPS monoclonal antibody and detected via Alexa-fluor 488 conjugated secondary antibody. Nuclei were visualized via DAPI staining (panels B and D). *Data shown are representative. Arrows show smaller vs. larger inclusion; magnification 60x*.

We further examined *C. trachomatis*-L2 infected hOE-WT and hOE-TLR3KO cells to quantitatively determine the impact of TLR3 deficiency on chlamydial inclusion development and size via multi-spectral imaging flow cytometry using the Amnis Image Stream X MKII. Figure 13 shows that cells deficient in TLR3 expression exhibited significantly increased levels of LPS within the chlamydial inclusion, based on the geometric mean fluorescence intensity differences (ΔMFI) between the two cell types. The fluorescence results demonstrating increased fluorescence intensity in the hOE-TLR3KO cells corroborates the IF data showing that the chlamydial inclusions were larger and presumably contained more chlamydial LPS during TLR3 deficiency. Further analyses of the Multi-spectral imaging flow cytometry data in IDEAS software revealed that the average *Chlamydia* inclusion diameter was calculated to be 15.3μm and 17.2μm in the hOE-WT and hOE-TLR3KO cells, respectively. Fig. S4 shows a representative image of the multi-spectral imaging flow cytometry, in which we scanned (in triplicate) 10000 cells each of the hOE-WT and hOE-TLR3KO cells that were either mock-infected or infected with *C. trachomatis*-L2. Collectively, immunofluorescence analyses and the imaging flow cytometry results using antibody specific for chlamydial LPS indicate that TLR3 deficiency in human oviduct epithelium leads to increased chlamydial inclusion size and more LPS within the inclusion.

**FIGURE 13.**
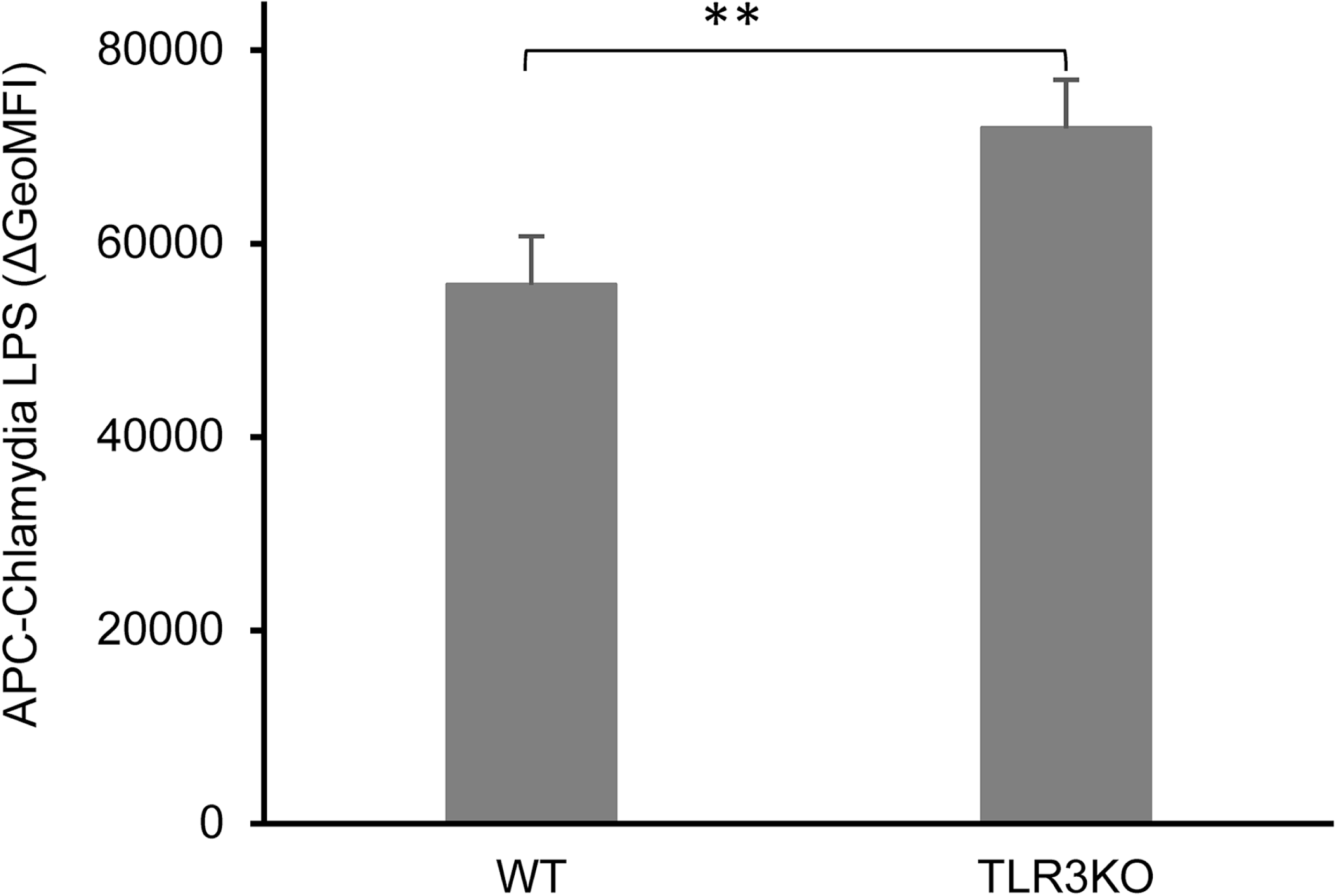
LPS levels within the chlamydial inclusions of infected WT and TLR3-deficient hOE cells. hOE-WT and hOE-TLR3KO cells were either mock treated or infected with *C. trachomatis*-L2 (in triplicate) at MOI of 5 IFU/ cell for 72hrs. *Chlamydia* LPS levels were determined in multispectral flow cytometric analyses of hOE cells stained using anti-*Chlamydia* LPS monoclonal antibody and allophycocyanin (APC) conjugated secondary antibody. APC-conjugated anti-IgG served as an isotype staining control. 10000 cells/events were processed in flow cytometry analysis. ΔMFI, Δ Geometric Mean Fluorescent Intensity. Data shown are representative of three independent experiments. Statistically significant differences are shown by asterisks (**, *p*<0.01).

To examine whether *Chlamydia* replication during TLR3-deficiency correlates with the increased inclusion size and LPS content within the inclusion, we next measured chlamydial replication in wild-type and TLR3-deficient hOE cells that were infected with 5 IFU/cell *C. trachomatis*-serovar D. As shown in Figure 14, *C. trachomatis* replication was greater at all time points in the hOE-TLR3KO cells compared to the wild-type, suggesting that stimulation of TLR3 by *C. trachomatis* results in an immune response that negatively affects *Chlamydia* growth. Interestingly, our data in Figure 14 show that the chlamydial replication in hOE-WT cells peaks at around 48hr post-infection before the recovery of infectious progeny begins to drop. In contrast, the recovery of infectious EBs in the *Chlamydia*-infected hOE-TLR3KO cells continued to rise past the 48hr time point (Fig. S5). Collectively, our data indicate that TLR3 plays a role in limiting *Chlamydia* replication in the HUMAN oviduct tissue and corroborates our previous studies in mice, and thus implicate TLR3 in a protective role against genital tract *Chlamydia* infections in both species.

**FIGURE 14.**
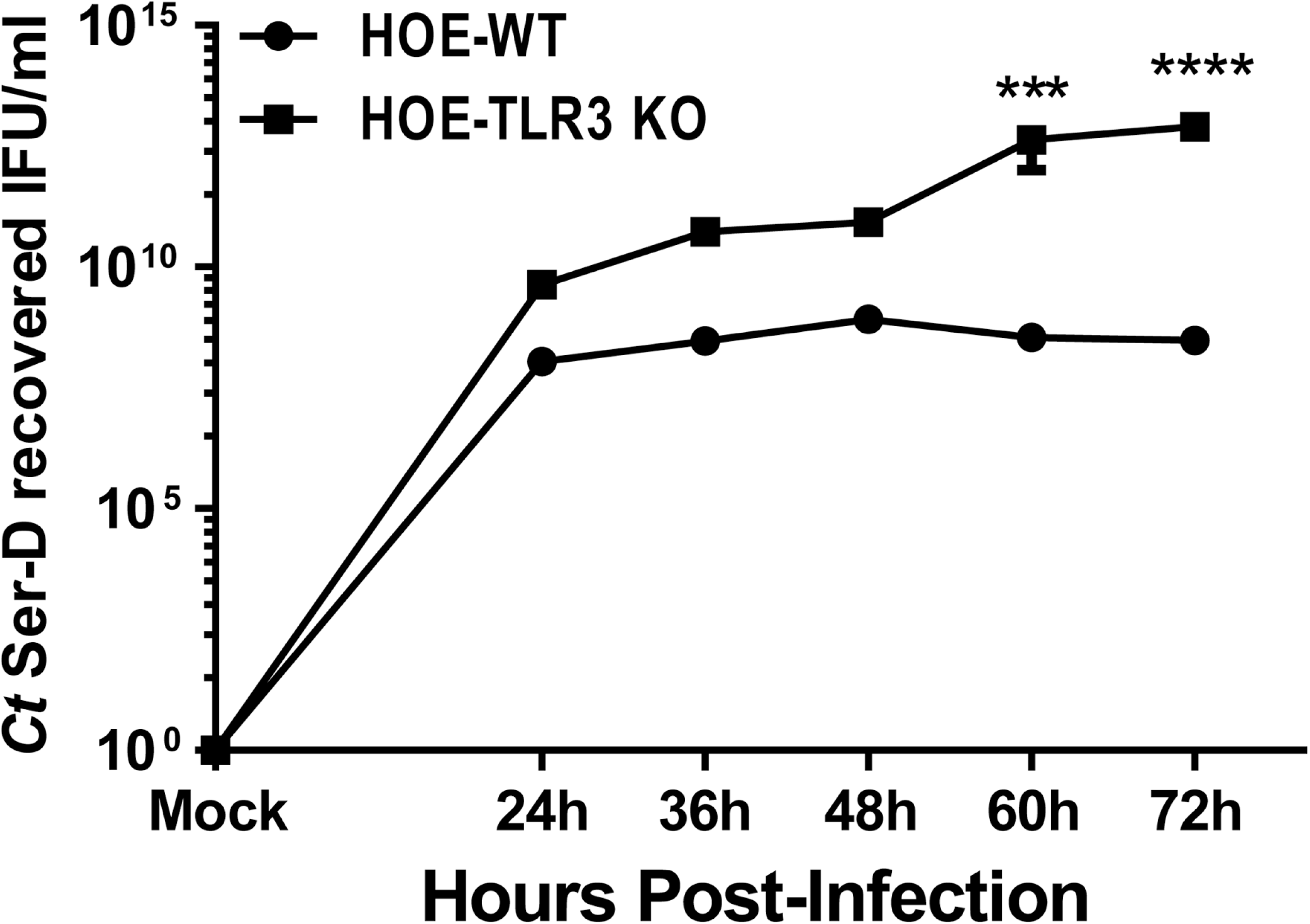
Measuring chlamydial replication in infected WT and TLR3-deficient hOE cells. hOE-WT and hOE-TLR3KO cells were either mock treated or infected with *C. trachomatis*-serovar D (in triplicate) at MOI of 5 IFU/ cell for 72hrs. The cells were disrupted in SPG buffer at the indicated time points as described in Materials and Methods and lysates were collected, sonicated, and titered on fresh Hela cell monolayers. The data presented are representative of three different experiments. Statistically significant differences are shown by asterisks (***, *p*<0.005; ****, *p*<0.001). IFU, inclusion forming units.

## DISCUSSION

Our research focuses on the impact of TLR3 signaling on the immune response to chlamydial infection in oviduct epithelium, and we were the first to demonstrate more severe genital tract pathogenesis in mice deficient in TLR3 confirming our hypothesis that TLR3 has a protective role in the immune responses to murine genital tract infections (32). In this investigation, our goal was to ascertain whether TLR3 had a similar protective role in the immune response to genital tract *Chlamydia* infection in humans and to more precisely delineate those immune responses in oviduct epithelial cells that contribute to the fibrosis and scarring that lead to reproductive sequelae in clinical disease. The OE-E6/E7 cells used in this report were derived from human Fallopian tubes and were immortalized by HPV 16 E6/E7 open reading frame (ORF) by retroviral expression (26). Although these are not primary oviduct epithelial cells, the cells are immortalized and offer the advantage of being able to be passaged in the laboratory, and they are close enough to primary oviduct epithelial cells that they can serve as an adequate representation of what we believe occurs during *in vivo Chlamydia*-OE cell interactions during natural genital tract infection.

Because the OE-E6/E7 cells express a functional TLR3, we first disrupted its gene expression and subsequent protein function using CRISPR to generate a TLR3-deficient version of the OE-E6/E7 cells, which could then be used to help delineate those immune responses to *Chlamydia* infection that are directly related to TLR3 function. We were able to generate several clones of the TLR3 deficient hOE cells and demonstrated loss of both TLR3 protein expression and its functional response to the TLR3 agonist poly (I:C). We also demonstrated a significant reduction in the chlamydial induced synthesis of IFN-β in hOE cells deficient in TLR3 function; however, the reduction was a bit more modest in the hOE-TLR3KO cells when compared to that of TLR3-deficient OE cells derived from TLR3KO mice (13). The differences that we observed in the reduction in the *Chlamydia*-induced IFN-β synthesis between the TLR3-deficient hOE and the murine OE cells are likely related to the fact that we were not able to completely disable TLR3 gene expression using CRISPR, whereas the functional TLR3 gene expression in the TLR3KO mouse is completely absent. Other possibilities to explain the more modest reduction in the *Chlamydia*-induced synthesis of IFN-β in the hOE cells could be more significant contributions of other pathways identified to enhance to the *Chlamydia*-induced type-I IFN response such as cyclic GMP–AMP (cGAMP) synthase (cGAS) and STING in the hOE cells (16, 66). The significance of the relatively higher level of IFN-β synthesis observed during *Chlamydia* infection in the TLR3-deficient hOE versus that of the TLR3-deficient murine OE cells is not yet known; however, we have shown that IFN-β can regulate the chlamydial-induced syntheses of a multitude of other inflammatory mediators (15).

As sentinels for invasion by microbial pathogens, epithelial cells lining the reproductive tract secrete cytokines and chemokines upon chlamydial infection that functions in various facets of innate immunity such as inflammation, lymphocyte recruitment, polarization, and genital tract scarring (24, 67). We have reported that TLR3 regulates the syntheses of a multitude of these factors both *in vitro* and *in vivo* in murine OE cells and mice, and concluded that TLR3 has regulatory function affecting multiple aspects of both the innate and adaptive immune response (13, 15, 32). We demonstrate in this report that TLR3 may have a similar role in human oviduct epithelial tissue and extrapolate our findings to speculate that TLR3 deficiency may have a significant role in outcomes of infections in humans. As an example, we show in Figures 5-7 that TLR3 deficiency in hOE cells leads to significant reductions in several factors associated with the acute inflammatory response such as IL-6, IL-6Rα, sIL-6Rβ (gp130) and IL-8. IL-6Rα and gp130 are the two chains that comprise the IL-6 receptor, which is a type I receptor for the pleiotropic cytokine IL-6. IL-6 represents a keystone cytokine in infection, cancer, and inflammation (68), and has been shown to have a significant role in both inhibiting *C. muridarum* infection in mice and exacerbating its pathogenicity in the mouse genital tract (69). IL-8 is a known as neutrophil chemotactic factor known to be induced early during *Chlamydia* infection and was thought to be associated with pre-term delivery complications in pregnant women infected with *C. trachomatis* (70).

Loss of TLR3 function in the hOE cells did not result in a global down-regulation in the *Chlamydia*-induced syntheses of all mediators that shape the immune response, nor was its impact limited to that of the acute inflammatory response. Our data also demonstrate that TLR3 deficiency leads to downregulation and *upregulation* in the chlamydial-induced syntheses of several cytokines and chemokines that have an impact at various phases of the adaptive immune response, which can potentially affect long-term outcomes of infection in humans and impact genital tract pathology. Figures 6 and 7 show that the TLR3-deficient hOE cells produced significantly higher levels of the leukocyte recruiting factor CCL-5 (71) and the type III interferons IL-28A and IL-29 (72) when compared to WT hOE cells. The type III IFNs are hypothesized to have a role in the polarization of the immune response to *Chlamydia* infection by inhibiting the production of IL-13, IL-4 and IL-5, and thereby promoting the development of protective Th1 immunity to infection (43). Our investigations into the role of TLR3 in the pathogenesis of *C. muridarum* infection in mice show that TLR3-deficiency leads to significantly altered CD4^+^/CD8^+^ T-cell ratios, and increased lymphocytic infiltration into the uterine horns and oviducts by day 21 post-infection (32). The observation of significantly higher levels of the type III interferons IL-28A and IL-29 in the TLR3-deficient hOE cells supports a hypothesis that TLR3 may have a role in polarization of the immune response in humans as well, and it would manifest itself by having an effect on the recruitment of lymphocytes into the female reproductive tract in women infected with *C. trachomatis*.

We recently reported that TLR3 deficiency resulted in an increased frequency and severity in pronounced chronic sequelae (such as lymphocytic endometritis and hydrosalpinx) during late stages of *C. muridarum* genital tract infections in mice (32). In this report, we show that the TLR3-deficient hOE cells were dysregulated in the *Chlamydia*-induced syntheses of several biomarkers associated with chronic inflammation, and are suggestive that chronic clinical outcomes would occur in higher frequency in humans lacking functional TLR3. Collectively, our data implicate TLR3 in having some impact on regulating the incidence and severity in outcomes of chronic inflammation caused during genital tract *Chlamydia* infections in humans. TNFα is involved in the systemic chronic inflammation that causes many of the clinical problems associated with autoimmune disorders such as rheumatoid arthritis, ankylosing spondylitis, inflammatory bowel disease, psoriasis, and refractory asthma. (73–75). These disorders are sometimes treated by using a TNF inhibitor, many of which that either mimic the TNFα receptor (TNF-R1) to bind to and block its activity or is an actual monoclonal antibody that binds TNFα (76). TLR3 deficiency in hOE cells showed defective syntheses of the soluble tumor necrosis factor receptor 1 (sTNF-R1) during *C. trachomatis* infection. sTNF-R1 binds to an inactivates TNFα, a major cytokine associated with scarring of oviduct tissue and severe genital tract sequelae in *C. muridarum* infected mice (44). This finding suggests that the diminished presence of sTNF-R1 in the TLR3 deficient cells would result in reduced effectiveness at inactivating *Chlamydia*-induced TNFα, and a subsequent increased incidence of TNFα-mediated scarring. In that same regard, dysregulation in the expression of matrix metalloproteinases (MMPs) is known to directly impact the severity of genital tract fibrosis and scarring in mice infected with *Chlamydia* (50–52). The *Chlamydia*-induced synthesis of MMP-1, MMP-2, MMP3, and MMP-10 was shown to be diminished in the TLR3-deficient hOE cells (Figure 9), suggesting that the normal physiological process of breaking down extracellular matrix proteins by the MMPs would be attenuated, and will likely result in increased fibrosis and scarring observed when the expression levels of certain MMPs are not sufficient (77, 78).

Our data showed that TLR3 deficiency in hOE cells significantly altered the *Chlamydia*-induced expression levels of biomarkers for chronic inflammation, persistence, and autoimmunity such as soluble CD163 (sCD163), Chitinase-3-like 1, Osteopontin (OPN), and Pentraxin-3 (58–60, 79–82). OPN was first identified in osteoclasts as an extracellular structural protein of bone and is known as an essential factor in bone remodeling (83). However, subsequent to the initial identification of OPN as a structural component of bone tissue, OPN has been shown to be expressed in a wide range of immune cells, including macrophages, neutrophils, dendritic cells, and various lymphocyte types, and is now known to have function in several aspects of host immunity (84). The exact role that OPN plays in the immune response to *Chlamydia* infection is poorly understood; however, recent studies have linked OPN to persistent inflammation and has hypothesized a role for OPN in cell transformation during persistent *Chlamydia* infection (85, 86). Our data showing significant reductions in OPN production in the TLR3-deficient hOE cells during *C. trachomatis* infection suggest that TLR3 deficiency may lead to increased incidences of persistent infections, and proposes a role for TLR3 in the prevention of long-term chronic inflammation and possible cellular transformation. A link between sCD163, Chitinase-3-like 1, Pentraxin-3 and *Chlamydia* infection has not yet been established; however, the dysregulation of these known biomarkers for chronic inflammation, persistent infection, and autoimmunity supports a role for TLR3 in limiting the clinical symptoms of chronic inflammation during infection.

Finally, we examined the possible impact that TLR3 deficiency in hOE cells may have on chlamydial replication and inclusion structure. We previously showed that *C. muridarum* replication was more robust, the inclusions were larger and more aberrantly shaped in TLR3-deficient murine OE cells, and that TLR3-deficient mice sustained significantly higher bacterial burdens than WT mice during early and mid-infection [(15, 32); Fig. S3]. We hypothesized from those studies that the overall negative effect on *C. muridarum* biology was largely due to the host-beneficial impact of TLR3-dependent IFN-β synthesis, and we showed that chlamydial replication in TLR3-deficient OE cells pre-treated with exogenous recombinant IFN-β was significantly diminished (15). Here, we corroborate the mouse studies by showing that the inclusions were substantially larger, and they were stained in a more punctate and diffuse pattern in TLR3-deficient hOE cells 36hrs after infection with *C. trachomatis*-serovar D. Imaging flow-cytometry revealed that TLR3-deficient hOE cells had higher levels of *Chlamydia* LPS within the chlamydial inclusion at 24hr post-infection when infected with *C. trachomatis*-L2 and are highly suggestive that the larger inclusions contain more bacterial particles. We consistently observed higher levels of *Chlamydia* replication in the TLR3-deficient hOE cells at all time-points between 24 and 72hrs post-infection (Figure 14 and Fig. S5) and that the differences were statistically significant after 48hs post-infection. Our data showing that TLR3 deficiency in hOE cells leads to the significant decreases in the chlamydial-induced synthesis of IFN-β implicates this cytokine in the control of *Chlamydia* infection in human genital tract epithelial cells as was observed in mice. However, our data in Figure 11 showed that TLR3-deficiency in hOE cells also resulted in drastic reductions in the chlamydial-induced secretion of lipocalin-2, which is an innate-immune protein that limits bacterial growth by sequestering iron-containing siderophores (87). Although it is unclear exactly which mechanism exerts the greatest impact on regulating chlamydial growth in hOE cells, our data are suggestive that there are likely redundant mechanisms that control *Chlamydia* replication that are disrupted during TLR3 deficiency.

Collectively, all of our previous investigations into the impact of TLR3 in the immune response to *Chlamydia* infection in mice have demonstrated a protective role for TLR3 in regards to genital tract pathology, and our data represents the first of such in regards to *Chlamydia* infection. Although the exact mechanism(s) that TLR3 invokes to elicit this protective immunity is unknown and needs further study, our preliminary investigations implicate TLR3 function in upregulating gene expression of other TLRs known to have impact on genital tract pathology (such as TLR2; (12)), through pathways involving IFN-β synthesis (15). In this report, we expanded our investigations into examining the impact of TLR3 in the immune response to *Chlamydia* infection in human oviduct epithelial cells, and our initial results thus far show a similar role for TLR3 in the protective immune response in humans. Studies are currently underway to further investigate the role of this enigmatic Toll-like receptor in human genital tract *Chlamydia* infection.

## ACKNOWLEDGMENTS

We thank Dr. David Nelson and his lab members for providing *C. trachomatis*-L2 strain, Serovar D, *Chlamydia* LPS antibody, for the use of the EVOS system, their thoughtful discussion, and their support. We also thank Dr. Cheikh Seye for access to the Wes™ Simple Western system. We thank the Multiplex Analysis Core at Indiana University’s Melvin and Bren Simon Cancer Center for providing support in analyzing samples and interpretation of data. This work was supported by NIH Grant AI104944 to W.A.D.

